# Anxiety Selectively Impairs Reward Learning Under Uncertainty, While N3 Sleep Recalibrates It

**DOI:** 10.64898/2026.01.01.696994

**Authors:** Rakshita Deshmukh, Arjun Ramakrishnan

## Abstract

Adapting to uncertain environments requires distinguishing stochastic variability from genuine environmental change, yet how anxiety and sleep shape this process remains unclear. Using a probabilistic reversal-learning task that simultaneously manipulated stochasticity and volatility, we show that high trait anxiety selectively impairs reward learning in stable but noisy environments, despite preserved sensitivity to true volatility. Computational modeling revealed that anxious individuals misattribute stochastic fluctuations to environmental change, resulting in elevated reward learning rates and impaired evidence accumulation. In a second experiment combining task performance with overnight sleep EEG, greater N3 sleep was associated with reduced learning rates and improved evidence accumulation. These findings identify a selective computational mechanism underlying anxiety-related learning deficits and suggest N3 sleep as a state-dependent pathway for recalibrating adaptive learning under uncertainty.

## Introduction

Adaptive behavior under uncertainty requires distinguishing stochasticity, random outcome variability under stable contingencies, from volatility, which reflects genuine environmental change. Normative learning models predict that learning rates should increase under volatility but decrease under high stochasticity to avoid overfitting noise (1-2). However, learning under uncertainty is also shaped by internal states such as anxiety, which is associated with intolerance of uncertainty and altered belief updating (3-5).

Trait anxiety, a stable vulnerability factor for anxiety disorders, is consistently linked to maladaptive learning and decision-making under uncertainty. Prior work has attributed these impairments to heightened learning from negative outcomes, reduced evidence accumulation, and disrupted medial prefrontal and anterior cingulate regulation, collectively inflating perceived environmental volatility (5-10). Anxiety has also been linked to aberrant salience attribution and learning deficits spanning reward and punishment domains (11,12). Critically, it remains unclear whether anxiety reflects a global hypersensitivity to environmental change, or instead a selective failure to discount stochastic outcome variability while preserving sensitivity to true volatility. Recent evidence suggests that some learners conflate stochasticity with volatility, increasing learning rates in noisy but stable environments (13), yet most studies have manipulated volatility in isolation, leaving unresolved how anxiety shapes learning when multiple sources of uncertainty must be inferred simultaneously.

Learning-rate regulation depends on neuromodulatory and prefrontal mechanisms that are strongly state-dependent, raising the possibility that physiological states such as sleep may selectively modulate uncertainty inference. Anxiety and sleep are tightly intertwined: sleep disruption exacerbates anxiety, while anxiety fragments sleep, jointly worsening outcomes (14,15). Sleep loss increases state anxiety and impairs prefrontal control, whereas non-rapid eye movement stage 3 (N3) sleep reduces anxiety and supports synaptic plasticity and learning (16–19). Whether N3 sleep can specifically normalize anxiety-related distortions in learning-rate adjustment under uncertainty, rather than exerting global effects on affect or performance, remains unknown.

Here, across two preregistered experiments (AsPredicted, I: https://aspredicted.org/cr8p7i.pdf; II: https://aspredicted.org/vy89xw.pdf), participants performed a probabilistic reversal-learning task in which volatility and stochasticity were simultaneously modulated. We show that high trait anxiety is associated with a selective failure to discount stochastic outcome variability, despite preserved sensitivity to true volatility, leading to impaired performance in stable yet noisy environments. Computational modeling revealed elevated reward learning rates alongside reduced evidence accumulation, contradicting accounts emphasizing exaggerated learning from negative feedback. In a second experiment combining task performance with overnight EEG, individual differences in N3 sleep predicted reduced morning state anxiety and selective normalization of learning in high trait-anxious individuals, reflected in attenuated reward learning rates and improved decision-making in noisy but stable environments. Together, these findings identify a computational phenotype linking anxiety to uncertainty misattribution and suggest N3 sleep as a state-dependent mechanism through which adaptive learning under uncertainty can be recalibrated.

## Results

Across two preregistered experiments (AsPredicted, I: https://aspredicted.org/cr8p7i.pdf; II: https://aspredicted.org/vy89xw.pdf), participants performed a three-option probabilistic reversal-learning task in which environmental uncertainty was independently manipulated along two orthogonal dimensions: stochasticity (reward probability) and volatility (rate of contingency change), allowing these factors to be dissociated within a single task.

In Experiment 1, a 3×2 within-subject design varied stochasticity (Low, Medium, High) and volatility (Slow, Fast). Fifty participants completed the task (N=50, 20 females, 22.32 ± 2.85 years), and trait anxiety was assessed using the STAI-Y (Fig 1A).

**Fig 1.**
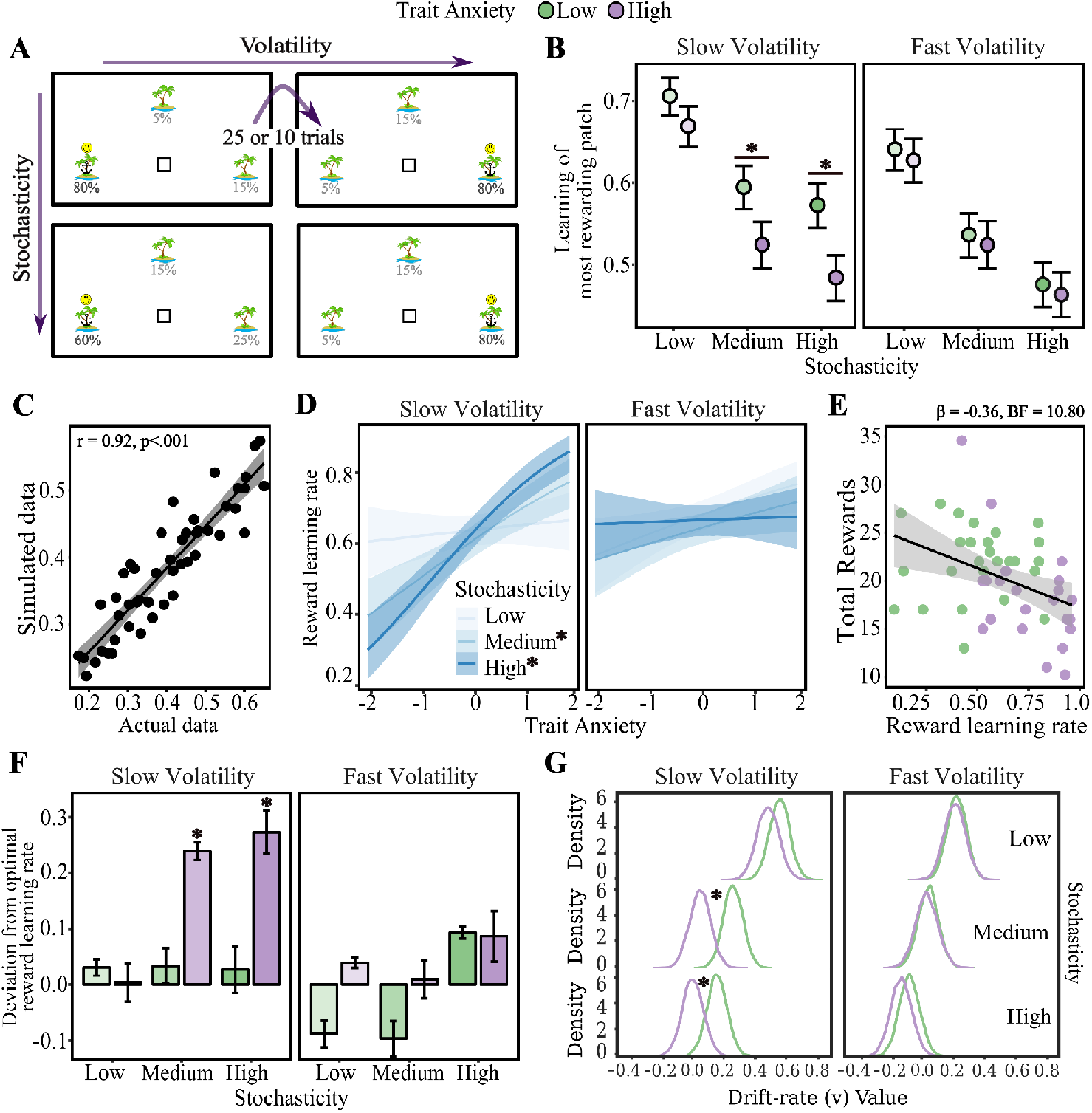
Trait anxiety impairs reward accumulation by driving suboptimally elevated reward learning rates under stochastic environments. Experimental design: A three-option probabilistic reversal-learning task with simultaneous stochasticity (Low, Medium, High with most-rewarding patch reward probability: 80%, 70%, 60%) and volatility (Slow or Fast, contingency changes every 25 or 10 trials in a 50 trial block, block order counterbalanced). On each trial, participants selected a patch from a central start position and received reward (smiley) or no-reward (sad) feedback to maximize earnings. Sample reward probabilities displayed on each patch, unbeknownst to the participants. **(B)** Posterior means (±SEM) of trial-wise learning of the most rewarding patch. As stochasticity and volatility increases, individuals with high trait anxiety (HTA) further lower learning in slow volatility but not fast volatility environments with this effect being largest in slow volatility-high stochasticity environments (Bayesian zero-one-inflated Beta Regression). **(C)** Parameter recovery: Average participant proportion of patch switches in simulated and actual data for hierarchical Bayesian reinforcement learning model showing significant positive correlation (*p*<.05, Pearson’s correlation), suggesting that the model captured the data well. **(D)** Posterior estimates of individual reward learning rates obtained from Hierarchical Bayesian reinforcement-learning modeling shows that, under slow volatility, higher trait anxiety is associated with increasing reward learning rates as stochasticity increases, with anxiety modeled as a continuous variable (Bayesian beta regression). **(E)** Participant environment-specific total rewards show higher reward learning rates are associated with significantly lower rewards in slow volatility-high stochasticity environment with HTA being clustered towards this regime (*BF*>1, Bayesian correlation). **(F)** Bar plots show the mean (± SEM) deviation from the optimal value (individual reward learning rate - optimal value) for each environment. HTA individuals significantly deviated from the optimal value in slow volatility high stochasticity environments (BF > 1; Bayesian one-sample *t*-tests), but not in fast volatility-high stochasticity environments. **(H)** Posterior distribution of drift-rates from Hierarchical Drift-Diffusion Modelling shows that while higher stochasticity and fast volatility lower drift-rates, HTA exhibits even lower drift-rates in slow volatility-high stochasticity environments, with values centered around zero (*q*<.05, comparing overlap proportions) but there is no difference between the two anxiety groups in fast volatility environments. *Note*: Asterisks (*) denote significant effects based on inferences from Bayesian statistics. For all regression models, reference levels were set as low stochasticity, slow volatility, and LTA.

In Experiment 2, we examined sleep-dependent modulation of learning using a 2×2×2 within-subject sleep design, with task performance before and after sleep under low versus high stochasticity and slow versus fast volatility. Forty participants completed the protocol (Fig 2A, N=40, 17 females, 22.75 ± 3.04 years). Anxiety was assessed pre- and post-sleep and overnight sleep was monitored using wearable EEG (see SM Methods).

**Fig 2.**
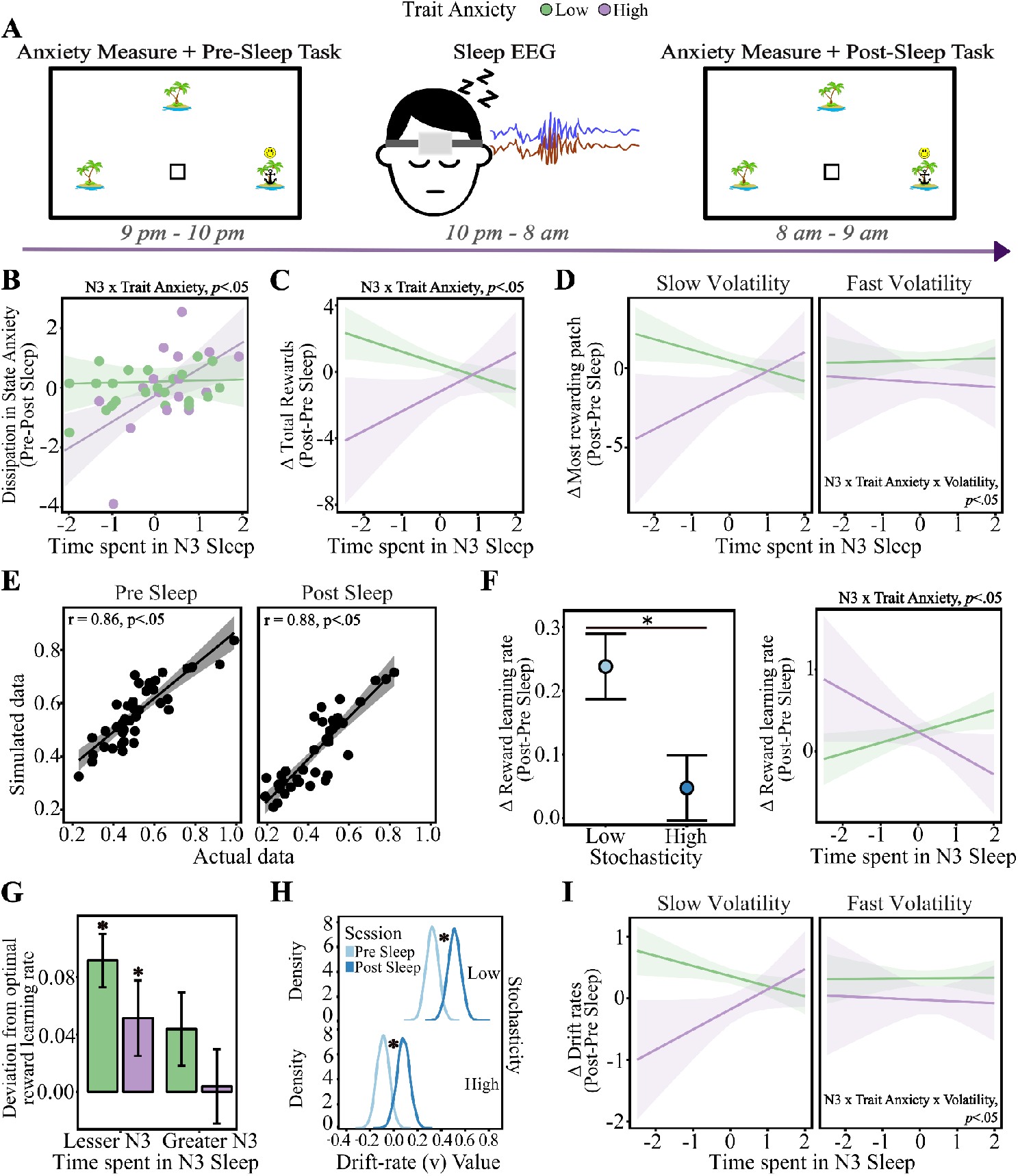
N3 sleep is associated with improved post-sleep performance in high trait anxiety (HTA) by reducing reward learning rates and enhancing patch valuation under slow volatility environments. **(A)** Experimental design: Participants completed a three-option probabilistic reversal-learning task, similar to Experiment 1, twice: pre- and post-sleep, with block order counterbalanced across sessions and participants. Task conditions manipulated stochasticity (low, high) and volatility (slow, fast), while anxiety was assessed using the STAI-Y scale before and after sleep. Participants slept in the laboratory, with sleep recorded using the Hypnodyne Zmax wearable EEG device. N3 sleep significantly interacts with trait anxiety in the dissipation of state anxiety (pre-post sleep state anxiety score), with higher values indicating positive post-sleep reductions in state anxiety; greater N3 sleep is associated with increasingly positive dissipation of morning state anxiety in HTA (*p*<.05, Linear Regression). **(C-D)** The benefit of N3 sleep for HTA extends to task performance, with greater N3 sleep associated with higher post-sleep rewards overall and improved identification of the most rewarding patch in slow-volatility environments (*p*<.05, Linear Mixed-Models). **(E)** Parameter recovery: Average participant proportion of patch switches in simulated and actual data per task session for hierarchical Bayesian reinforcement learning model showing significant positive correlation (*p*<.05, Pearson’s correlation), suggesting that the model captured the data well. **(F)** Individual level reward learning rates from the Hierarchical Bayesian Reinforcement Learning model shows that post-sleep reward learning rate decreases in high stochasticity (left), with greater N3 sleep reducing HTA’s overall reward learning rates (right) (*p*<.05, Linear Mixed-Models). **(G)** Bar plots show the mean (± SEM) deviation from the optimal value (individual reward learning rate - optimal value) for post-sleep session. Greater N3 sleep is associated with reduced deviation from the optimal learning rate, with no significant deviation observed in either HTA or LTA, whereas lesser N3 sleep (categorical mean split) shows significant deviation in both groups, indicating that N3 sleep stabilizes post-sleep reward learning, particularly in HTA (*BF*>1, Bayesian one-sample t-tests). **(H)** Posterior distribution of drift-rates from Hierarchical Drift-Diffusion Modelling shows post-sleep drift rates increase across environments (*q*<.05, comparing posterior proportions). **(I)** At the individual level, greater N3 sleep increases higher post-sleep drift rates in HTA, particularly in slow-volatility environments (*p*<.05, Linear Mixed-Models). Fast volatility environments remain unaffected by anxiety or sleep. *Note*: Asterisks (*) denote significant effects based on frequentist statistics, whereas * indicates inferences from Bayesian statistics. Variables were normalized; high and low trait anxiety reflect a mean split for plotting, with continuous values used for statistical analyses. For all regression models, reference levels were set as low stochasticity and slow volatility.

### Trait anxiety impairs reward accumulation in stochastic environments

We tested whether increasing environmental uncertainty reduced reward accumulation, and whether this effect was exacerbated in individuals with high trait anxiety (HTA; STAI-T ≥ 45) relative to low trait anxiety (LTA). Reward accumulation decreased with stochasticity (non-parametric ANOVA; *F*_2,276_=74.16, *p*<.001, η^2^ _p_ =0.35) and volatility (*F*_1,276_=7.69, *p*<.05, η^2^ _p_ =0.03), with graded effect of stochasticity (Low > Medium > High; all *p*<.05; Table S1) and volatility levels (*p*<.05).

Trait anxiety further modulated reward accumulation with a significant main effect (*F*_1,276_=4.08, *p*<.05, η^2^_p_ =0.02) and Trait Anxiety x Volatility interaction (*F*_2,276_=4.87, *p*<.05, η^2^ _p_ =0.02). This interaction was driven by reward accumulation by HTA individuals under slow volatility (simple effects; Slow: *F*_1,142_=5.44, *p*<.05,η^2^_p_ =0.04), with no anxiety-related differences under fast volatility (Fast: *F*_1,142_=0.01, *p*=.9,η^2^_p_=0.00). Consistent with this pattern, HTA participants accumulated fewer rewards than LTA under slow volatility (Pairwise t-tests:*t*_128_=−2.3, 95%62 [−CI.04, −4.69], *p*<.05, *d*=−0.39).

Environment-specific analyses revealed that this anxiety-related deficit was selective to high-stochasticity, slow-volatility conditions, where HTA participants earned significantly fewer rewards than LTA counterparts (*t*_37_=−2.61, 95%CI [−6.32, −0.7], *p*<.05, *d*=−0.77), with no group differences in other environments (Table S2).

To examine learning dynamics underlying these effects, we modeled trial-by-trial identification of the most rewarding option using Bayesian Beta regression (See SM Methods). Learning accuracy decreased with increasing stochasticity and volatility (Fig 1B). Critically, trait anxiety selectively impaired learning under slow volatility, in medium and high stochasticity environments (Fig 1B, left), with no anxiety-related differences under fast volatility (Fig. 1B, right). HTA was associated with reduced learning of the most-rewarding patch (β=−0.17, 95%CI [−0.32, −0.01]), with additional negative effects of medium (β=−0.12, 95%CI [−0.18, −0.05]) and high stochasticity (β=−0.19, 95%CI [−0.25, −0.13]; Table S3).

Together, these findings indicate that high trait anxiety selectively impairs performance in stable environments characterized by high stochasticity.

### Trait anxiety is associated with maladaptively elevated reward learning rates under stochastic environments

We preregistered the hypothesis (I) that HTA individuals would be associated with elevated punishment relative to reward learning rates under high uncertainty. However, given the reduced performance under slow volatility, we first tested whether heightened punishment learning accounted for impaired learning in stable environments. Contrary to this prediction, HTA was associated with higher reward learning rates under slow volatility, selectively in medium and high stochasticity environments.

To estimate learning rates, we fit a 5-parameter hierarchical Bayesian reinforcement-learning model with separate learning rates for reward and punishment (see SM Methods). The model provided a good account of behavior (Fig. 1C). Under slow volatility, HTA individuals exhibited elevated reward learning rates in medium (95%HDI [0.11, 0.51]) and high stochasticity conditions (95%HDI [0.03, 0.39]), Tables S4-S5). This effect was replicated at the individual-level using a Bayesian Beta regression (Medium: β=−0.41, 95%CI [−0.73, −0.10]; High: β=−0.83, 95%CI [−1.16, −0.5], Tables S6-S7), and was robust when trait anxiety was modeled as continuous (Fig 1D left, increasing from left to right) or categorical variable (Fig S1). Notably, this finding runs counter to normative expectations, which predict lower learning rates in stable environments as stochasticity increases. However, under high volatility, reward learning rates did not vary as a function of trait anxiety (Fig 1D, right).

We next tested whether elevated learning rates were maladaptive. Higher reward learnings rates were associated with lower reward accumulation in the slow-volatility, high-stochasticity condition (Bayesian correlation: β=−0.36, 95%CI [−0.57, −0.09], BF=10.80) with HTA individuals clustering in this suboptimal regime (Fig 1E). Punishment learning rates were not associated with reward accumulation, indicating that elevated reward learning rather than punishment learning drove the performance deficit (Table S8).

To benchmark learning behavior against normative expectations, we simulated optimal learning rate values for each task environment (see SM Methods). Normative reward learning rates decreased with higher stochasticity and increased with faster volatility (Fig S2). Relative to these benchmarks, HTA participants showed a selective deviation from optimality in slow volatility, medium stochasticity (Bayesian one-sample t-tests: β=0.24, 95%CI [0.20, 0.27], BF=1.04e+10) and high stochasticity environments (β=0.26, 95%CI [0.18, 0.35], BF=33200, Fig. 1F, left), but not in fast volatility environments (Fig 1F, right, (Medium: β=0.01, 95%CI [−0.05, 0.07], BF=0.23; High: β=0.08, 95%CI [−0.02, 0.17], BF=1.04).

Punishment learning rates for both HTA and LTA participants remained below optimal and did not differ substantially under slow volatility, indicating that punishment learning did not contribute to reward maximization in this task (SM Text).

Finally, we tested whether suboptimal learning reflected impaired valuation of the most rewarding option. Using a hierarchical drift-diffusion model, we found that under slow volatility and high stochasticity, HTA participants exhibited reduced drift rates, centered near zero relative to LTA participant (Fig 1G, Hierarchical Drift-Diffusion Model, High: 95.24%, *q*<.05, Medium: 98.97%, *q*<.05, Table S9, S10), along with higher decision boundaries in this environment (95.08%, *q*<.05). Bias and non-decision time did not differ between groups.

This anxiety-related impairment was selective to slow-volatility environments and was absent under fast volatility, where neither learning rates nor evidence accumulation differed between anxiety groups (Fig 1G).

Collectively, these findings indicate that high trait anxiety is associated with maladaptively elevated reward learning rates in stable yet noisy environments, reflecting a misinterpretation of stochastic outcome variability as genuine environmental change.

### N3 sleep lowers morning anxiety and improves performance in high trait anxious individuals

In experiment 1, we confirmed a bidirectional relationship between anxiety and sleep: higher trait anxiety was associated with poorer subjective sleep quality (Sleep Condition Indicator; Spearman’s r(46) = −0.41, p <.05), consistent with prior reports linking anxiety to sleep disruption (15).

For Experiment 2, we preregistered the hypotheses (II H1a, H1c iii) that greater N3 sleep would predict lower next-day state anxiety based on previous findings (16), and that LTA participants would show greater N3 sleep and larger overnight anxiety reduction than HTA individuals. Across participants, greater N3 sleep was associated with larger evening-to-morning reduction in state anxiety, with no comparable associations for N1, N2, or REM sleep. (Pearson’s correlation, r(38)=0.36, *p*<.05, table S12). Linear regression revealed significant main effect of N3 sleep (β=0.46, 95%CI [0.15, 0.77], t_32_=3.01, *p*<.05) and an interaction with trait anxiety (β=0.43, 95%CI [0.06, 0.79], t_32_=2.39, *p*<.05, Table S13).

Contrary to our hypothesis, HTA participants showed the greatest anxiolytic benefit from N3 sleep (Fig 2B, in purple, increasing from left to right).

We next examined whether sleep-related anxiety reduction translated into improved task performance. We preregistered that poor sleep would impair performance, particularly in HTA individuals (II H2a, H2b). Across participants, post-sleep reward accumulation was higher than pre-sleep (ANOVA: *F*_1,312_=8.907, *p*<.05,η^2^_p_=0.03; Post-hoc t-test:t_159_=3.33, *p*<.05, *d*=0.26, 95%CI [0.68, 2.66]), while stochasticity and volatility reduced rewards across both sessions (Table S14). Critically, greater N3 sleep predicted higher post-sleep reward accumulation in HTA participants (Fig 2C, Linear Mixed-Model, β=0.46, 95%CI [0.03, 0.88], t_133_=2.13, p<.05, Table S15, S16, SM text). This benefit was driven by improved ability to identify the most rewarding patch under slow volatility (Fig 2D left, in purple, increasing from left to right; Linear Mixed-Model: β=−0.49, 95%CI [−0.92, −0.07], t_120_=−2.28, *p*<.05, Table S17), but not under fast volatility (Fig. 2D, right).

Together, these results indicate that greater N3 sleep is associated with reduced morning anxiety and selective restoration of HTA individual’s performance especially in stable but noisy environments.

### N3 sleep ameliorates high trait anxious individual’s maladaptive reward learning rates

We preregistered that poor sleep would exacerbate punishment learning in high trait-anxious (HTA) individuals under high uncertainty, whereas good sleep would enhance reward learning in low trait-anxious (LTA) individuals (II H2c & H2d). However, Experiment 1 instead revealed that HTA individuals exhibited *elevated reward learning rates* in slow-volatility, high-stochasticity environments, and Experiment 2 showed that N3 sleep reduced morning state anxiety and improved performance in HTA participants. These findings motivated tests of whether N3 sleep selectively modulates reward learning in environments where HTA show maladaptive updating.

Reward learning rates were estimated using a hierarchical Bayesian reinforcement-learning model, which provided a good account of behavior pre- and post-sleep (Fig. 2E). Across participants, reward learning rates decreased from pre-to post-sleep under high stochasticity, consistent with normative learning-rate adjustment. Importantly, greater N3 sleep was associated with reduced post-sleep reward learning rates in HTA individuals across all environments (Fig 2F, left, Linear Mixed Model: Stochasticity: β=−0.92, 95%CI [−1.16, −0.69], *p*<.05; N3 x Trait Anxiety: β=−0.45, 95%CI [−0.86, −0.03], *p*<.05, Table S18-S20). Focusing specifically on slow volatility-high stochasticity environment where HTA showed elevated learning rates in Experiment 1, greater N3 sleep significantly reduced post-sleep reward learning rates in HTA individuals (Fig 2F, right, in purple, decreasing from left to right; Linear regression: β=−0.09, 95%CI [−0.17, −0.01], *p*<.05).

To assess whether N3 sleep shifted learning rates from suboptimal toward optimal values, we compared pre- and post-sleep learning rates to simulated optimal values, before and after sleep. In the pre-sleep session both HTA and LTA groups significantly deviated from the optimal reward learning rate (Bayesian one-sample t-tests: HTA: β=−0.07, 95%CI [−0.13, −0.02], BF=5.15; LTA: β=−0.07, 95%CI [−0.14, −0.04], BF=37.74). But, post-sleep, greater N3 sleep was associated with learning rates that were near optimal values for both groups, whereas less N3 sleep was associated with significant deviations from optimal values (Fig 2G, Bayesian one-sample t-tests: HTA-More N3: β=0.004, 95%CI [−0.06, 0.07], BF=0.30; LTA-More N3: β=0.04, 95%CI [−0.02, 0.10], BF=0.62; HTA-Less N3: β=0.04, 95%CI [0.01, 0.08], BF=3.22; LTA-Less N3: β=0.08, 95%CI [0.02, 0.14], BF=11.14). Thus, greater N3 sleep shifted HTA’s elevated reward learning rates towards optimal values.

Finally, we tested whether N3 sleep optimized reward learning in HTA by improving valuation of the most rewarding patch. Post-sleep drift rates were significantly higher than pre-sleep across all environments (Fig 2H, Hierarchical Drift-Difussion Model, Low-Slow=99.18%, *q*<.05; Low-Fast=99.93%, *q*<.05; High-Slow=98.63%, *q*<.05; High-Fast=97.28%, *q*<.05). Moreover, greater N3 sleep significantly interacted with trait anxiety overall and specifically under slow volatility environments in increasing post-sleep drift-rates in HTA individuals (Fig 2I, in purple, increasing from left to right; Linear Mixed Model: N3 x Trait Anxiety: β=0.48, 95%CI [0.03, 0.92], *p*<.05; N3 x Trait Anxiety x Volatility: β=−0.51, 95%CI [−0.93, −0.08], *p*<.05, Table 21-22). These changes indicate more efficient identification of the most rewarding patch by HTA individuals following sleep.

Together, these findings show that N3 sleep selectively ameliorates maladaptive reward learning in high trait anxiety by shifting learning rates toward optimal values and improving value-based decision-making.

## Discussion

Although anxiety-related impairments in learning under uncertainty are well documented (20, 21), the computational source of these deficits remain unclear. By jointly manipulating environmental stochasticity and volatility, we observed a selective performance deficit that arises due to misinterpretation of stochastic reward fluctuations as genuine environmental change in HTA individuals. While they remained sensitive to true environmental volatility (22), they failed to appropriately discount stochastic outcome variability. As a result, they exhibited elevated reward learning rates and slower evidence accumulation in stable yet noisy environments, leading to premature belief updating (23). Crucially, N3 sleep was associated with reduction in anxiety-related impairments, and recalibration of learning and post-sleep decision-making.

Computational modeling revealed that while high learning rates are adaptive in volatile environments, anxious individuals maintained elevated reward learning rates even in stable but noisy conditions, paradoxically increasing learning under stochasticity (13), failing to distinguish signal from noise, overestimating volatility (7, 24, 25). Anxious individuals treated random outcomes as informative environmental shifts, specifically in slow-volatility, high-stochasticity contexts. Consistent with prior links between anxiety and impaired decision-making under cognitive demand (9, 26-28), these effects were accompanied by reduced drift rates, indicating inefficient accumulation of reward evidence (29). Notably, the observed impairments were reward-driven rather than punishment-driven, challenging dominant accounts that attribute anxiety-related learning impairments primarily to enhanced punishment sensitivity or threat learning. Even in the absence of explicit penalties, noisy rewards appear to overwhelm accurate inference of environmental statistics in anxious individuals. This interpretation is consistent with evidence that anxiety amplifies sensitivity to salient monetary incentives under uncertainty (4, 11, 30), or whichever outcome appears most informative (12, 31).

Previously N3 sleep has been associated with reduction in morning state anxiety (16), which held true in our data as well. However, we also observed that HTA individuals benefited the most from N3 sleep. This benefit also extended to improved post-sleep learning in HTA participants, yielding higher rewards and more accurate identification of the optimal option. While N3 sleep supports optimal learning across domains, including motor learning and cognitive flexibility (19, 32, 33), in our task N3 sleep was associated with selective recalibration of learning rate and evidence accumulation, restoring the computational boundary between stochastic noise and environmental change disrupted by anxiety. While typical sleep deprivation studies have demonstrated exaggerated reward and threat reactivity and reduced sensitivity to environmental volatility, impairing belief updating (34-36), we demonstrate that sufficient N3 sleep is associated with regulation of volatility misestimation by improving the decision process.

Several considerations help define the scope of these findings. Because N3 sleep was not experimentally manipulated, its restorative effects remain partially correlational, motivating future causal tests using targeted modulation of N3 sleep. Furthermore, unmeasured circadian and chronotype factors may have influenced pre-versus post-sleep performance, highlighting the importance of jointly considering sleep and circadian contributions to learning under uncertainty. Notably, the observed effects were selective to environments requiring appropriate discounting of stochastic noise, arguing against nonspecific arousal or vigilance accounts.

Overall, these findings demonstrate that high trait anxiety promotes a maladaptive misestimation of stochastic noise as volatility, leading to elevated reward learning and inefficient evidence accumulation even in stable environments. In contrast, N3 sleep recalibrates these computations, normalizing reward learning rates and accelerating evidence accumulation. Because stochasticity-volatility trade-offs are fundamental to many real-world learning problems, these findings may generalize beyond the specific task, highlighting sleep as a critical modulator of how anxiety impacts adaptive learning and decision-making under uncertainty.

## Acknowledgments

This work was supported by the Science and Engineering Research Board start-up research grant SRG/2020/001723 (AR), and DBT Wellcome Trust Intermediate Fellowship IA/I/20/2/505204 (AR). We gratefully acknowledge the IISc-Pratiksha Trust’s Brain, Computer and Data Science Initiative for awarding a Senior Student Research Associateship to RD.

## Author contributions

Conceptualization: RD and AR; Data curation: RD; Formal Analysis: RD; Funding acquisition: AR; Investigation: RD and AR; Methodology: RD and AR; Software: RD; Resources: AR; Project administration: AR; Supervision: AR; Validation: RD and AR; Visualization: RD; Writing – original draft: RD; Writing – review & editing: RD and AR

