## Supplementary Materials for "Anxiety Selectively Impairs Reward Learning Under Uncertainty, While N3 Sleep Recalibrates It"

### Materials and Methods

Participants

Ninety young adults were recruited from the campus community of the Indian Institute of Technology Kanpur via email advertisements. All participants provided written informed consent. Ethics approval was obtained from the Institute Ethics Committee at IIT Kanpur (IITK/IEC/2020-21/II/10). Recruitment and participation for both experiments followed the exclusion criteria described below. The first behavioral experiment (N=50, 20 females, 22.32 ± 2.85 years) consisted of informed consent, questionnaire completion, and task performance, and was conducted in the Decision Lab at IIT Kanpur. The session lasted approximately 25-30 minutes. From this, 2 participants were excluded due to low alertness and extreme sleepiness, as assessed using the Stanford Sleepiness Scale (SSS). Thus, the final number of participants included in the analysis of the first experiment was 48 (19 females, 22.44 ± 2.83 years). The second experiment (N=40, 17 females, 22.75 ± 3.04 years) employed a repeated-measures design with overnight sleep recording in the Sleep Lab at IIT Kanpur. Participants arrived at the laboratory at 21:00, provided consent, and completed pre-sleep questionnaires assessing sleep and anxiety. Participants completed the behavioral task before sleep, after which sleep was recorded overnight using a wearable EEG system. The laboratory environment was temperature-controlled (26 °C). Participants were permitted to bring personal items to enhance comfort; pillows and disposable bedding were provided. The following morning, approximately 08:00 (±45 min), participants repeated the behavioral task and completed post-sleep anxiety questionnaires. For both experiments, participants received monetary compensation consisting of a fixed base amount and a performance-based bonus.

Pre-registration, power analysis and exclusion criteria

Hypotheses, planned sample sizes and exclusion criteria were preregistered for Experiment 1 (#123106) and Experiment 2 (#128319) on the AsPredicted platform. The sample size for Experiment 1 was determined using a power analysis based on pilot data, conducted with the *simr* (36) package in R, indicating that 40-50 participants were required to achieve 80% power. A target sample of 50 participants was therefore selected to allow for exclusions. For Experiment 2, prior sleep studies (16, 19, 38) informed sample size expectations (14-32 participants). Data were collected from 40 participants who met all inclusion criteria, exceeding minimum requirements. Across both experiments, participants were excluded if they reported an SSS score ≥ 3 or failed to sample from all three choice options during the task. For Experiment 2, additional exclusion criteria included age outside 18-35 years, history of sleep or neurological disorders, Axis I psychiatric disorders, substance abuse, closed head injury, or use of medications affecting sleep or cognition. Participants were instructed to abstain from alcohol, drugs, and caffeine on the day of testing aligning with standard sleep study exclusion criteria (16). The pre-registration can be accessed at [https://aspredicted.org/cr8p7i.pdf; https://aspredicted.org/vy89xw.pdf]. Any deviations are limited to the results and their interpretation; the methods remain unchanged.

Questionnaires

1. *Stanford Sleepiness Scale (SSS)*: Momentary alertness was assessed using the Stanford Sleepiness Scale (39), a single-item measure ranging from 1 (very alert) to 7 (very

sleepy). The SSS was administered prior to task performance to screen for low alertness. Participants reporting scores ≥3 were excluded from further participation. The SSS was not included in subsequent analyses.

2. *State-Trait Anxiety Inventory (STAI-Y)*: Trait and state anxiety were assessed using the State-Trait Anxiety Inventory (40), a 40-item self-report questionnaire comprising 20 items each for state (STAI-S) and trait (STAI-T) anxiety. In Experiment 1, state anxiety scores ranged from 20 to 60 (34 ± 9), and trait anxiety from 26 to 60 (44 ± 8.8). For analysis, participants were categorized as high trait anxious (HTA; STAI-T ≥ 45) or low trait anxious (LTA), consistent with prior literature (10, 41, 42). K-means clustering independently identified two clusters with a boundary at STAI-T=45, supporting this threshold. In Experiment 2, trait anxiety scores ranged from 30 to 72 (44.35 ± 10.01). State anxiety scores ranged from 21 to 60 at pre-sleep (33.45 ± 8.99) and from 20 to 63 at post-sleep (32.43 ± 9.62). Trait anxiety measured at the beginning of the experiment was used for all analyses. Dissipation in state anxiety was quantified as the difference between pre-sleep and post-sleep scores.
3. *Sleep Condition Indicator (SCI)*: Subjective sleep quality was assessed using the Sleep Condition Indicator (43), an 8-item questionnaire with scores ranging from 0 to 32, where higher scores indicate better sleep quality. Across both experiments, SCI scores ranged from 5 to 32 (22.98 ± 5.9).
4. *Post-sleep Questionnaires*: In Experiment 2, participants completed a post-sleep questionnaire adapted from prior studies (43, 44) to assess sleep experience and comfort. Participants reported sleep onset latency, number of nocturnal awakenings, perceived sleep duration, dream occurrence and valence, and rated sleep quality, morning sleepiness, and subjective refreshment on 7-point Likert scales. Comfort with the wearable EEG device was also rated. On average, participants reported a sleep onset latency of 35.0 min (± 24.88), 1.85 awakenings per night (± 1.23), and a perceived sleep duration of 7.02 h (± 1.47). Seventeen participants reported no dreams, sixteen reported pleasant dreams, five reported unpleasant dreams, and two reported dreams related to the experiment. Mean ratings indicated good sleep quality (5.15 ± 1.05), low morning sleepiness (2.63 ± 1.53), high refreshment (5.25 ± 1.25), and good device comfort (5.23 ± 1.54).

<u>Behavioral Task and Experimental Design</u>

Participants performed a three-option probabilistic reversal-learning task framed as a fishing game. Participants were instructed to choose among three fishing locations ("patches") on each trial, with each location delivering a catch with a fixed probability. Feedback was presented visually as a smiley emoji for reward and a sad emoji for no reward. Participants were informed that some fishing locations were generally more rewarding than others, but that reward contingencies could change over time from fishes swimming across the patches. They were not informed about the exact reward probabilities or reversal structure. This cover story was identical across all task conditions and experiments. The most rewarding patch delivered rewards with probabilities of 80%, 70%, or 60% depending on the stochasticity condition, while the remaining two patches had reciprocally lower probabilities (15% and 5%, 20% and 10%, or 25% and 15%, respectively). Reward contingencies reversed after a fixed number of trials determined by the volatility condition (after 10 or 25 trials in a 50 trial block). There was no time limit for responding. All task instructions were presented in writing on the computer screen, and participants completed five practice trials prior to the main task. Patch probabilities were randomized across participants, and within the session.

1. *Experiment 1* used a 3×2 within-subjects design with three levels of stochasticity (80%, 70%, 60%) and two levels of volatility (fast, slow). After completing questionnaires, participants completed six blocks (80fast, 80slow, 70fast, 70slow, 60fast, 60slow), each comprising 50 trials (300 trials total). Block order was counterbalanced across participants, with a 30-s break between blocks. The session lasted approximately 25-30 min. Participants received a fixed base payment of INR 30 with a performance-dependent bonus of up to INR 50.
2. *Experiment 2* employed a repeated-measures pre-sleep/post-sleep design. In each session, after completing questionnaires, participants completed a 2×2 within-subjects task with two levels of stochasticity (80%, 60%) and two levels of volatility (fast, slow). Each session comprised four blocks (80fast, 80slow, 60fast, 60slow), each containing 50 trials, yielding 400 trials across both sessions. Block order was counterbalanced across participants and within sessions. Each session lasted approximately 15 min and was separated by overnight sleep, which was recorded using a wearable EEG device. Participants received a fixed base payment of INR 350 for overnight participation, with an additional performance-dependent bonus of up to INR 100.

Sleep EEG recording and preprocessing

Sleep was recorded using the Hypnodyne ZMax, a portable EEG system validated against standard polysomnography (PSG) for whole-night sleep monitoring, with reliable capture of sleep-relevant EEG frequencies (0.3–30 Hz) and no reported adverse effects on sleep or mood (45, 46). The ZMax headband recorded EEG and eye-movement signals from two frontal bipolar channels (F7–FPz and F8–FPz), referenced to FPz, and sampled at 256 Hz. Following completion of the pre-sleep task, participants' foreheads were cleaned, a new disposable electrode patch was applied, and the headband was positioned centrally on the forehead and secured with an adjustable strap. Wireless recording was initiated using HDRecorder software, and signal quality was verified through a brief calibration procedure involving head movements and eye blinks. Participants wore the device continuously overnight. In the morning, recording was stopped, the device was removed, and participants completed post-sleep questionnaires and tasks.

Sleep was continuously recorded overnight for all participants with no data loss or disconnections. Sleep staging was performed in 30-s epochs according to American Academy of Sleep Medicine criteria using the YASA Python package,  an open-source Python-based package. YASA (Yet Another Spindle Algorithm) is trained on diverse datasets and demonstrates high accuracy comparable to human interrater agreement across sleep stages, ensuring reliable and generalizable sleep-stage scoring that is unaffected by body composition or participant sex and avoids systematic overestimation or underestimation of specific sleep stages (47). Sleep onset latency (SOL), total sleep time (TST), latency to each stage, and time spent in each sleep stage were computed separately for the left and right EEG channels. Raw EEG data were downsampled to 100 Hz and bandpass filtered between 0.3-45 Hz prior to spectral analysis. Power within standard frequency bands: delta (0.5–4 Hz), theta (4–8 Hz), alpha (8–12 Hz), sigma (12–16 Hz), beta (16–30 Hz), and gamma (30–40 Hz) was computed for each sleep stage and channel. Consistent with preregistration, only delta band power during N3 sleep was retained for subsequent analyses. Sleep spindle and slow-wave activity was also calculated using standard detection thresholds but were analyzed in this study. To assess concordance between EEG channels, preregistered sleep variables (TST, N3 duration, and SOL) were compared between left and right electrodes using Wilcoxon rank-sum tests. No significant differences were observed across any measure (TST: W=814, $p$=.9; N3: W=848, $p$=.7; SOL: W=731.5, $p$=.5); therefore, all analyses used data from the left EEG channel. Preregistered EEG-derived variables included time spent in N1, N2, N3, and REM sleep, SOL, TST, REM latency, number of REM episodes, and delta band power during N3 sleep. All variables were z-scored prior to statistical analysis. Summary sleep statistics are reported in Table S23.

<u>Behavioural Data Preprocessing and Statistical Analysis</u>

Raw behavioral data were cleaned and preprocessed in Python (v3.12.7). For each trial, reaction times and rewards earned were extracted. For each block, we computed whether participants identified the most rewarding patch (1 if they did, else 0) and a trial-wise cumulative learning score reflecting the proportion of choices directed toward the most rewarding patch relative to all prior choices. All statistical analyses were conducted in R (v4.4.2). Descriptive statistics, ANOVAs, and t-tests were performed using *rstatix* (48). Linear regressions, linear mixed-effects models, and mixed logistic models were fitted using *lmerTest* and *brms*, with model assumptions assessed using *performance* and visualizations generated using *ggplot2*, *sjPlot* and *interactions* (49-53). For all mixed models, reference levels were low stochasticity, slow volatility, and low trait anxiety. Trait anxiety was z-scored when modeled as a continuous predictor. Dissipation in state anxiety was defined as the evening-to-morning change in STAI-S scores (pre-post state anxiety), with positive values indicating reduced morning anxiety. Identification of the most rewarding patch was quantified as the proportion of correct choices per block. All survey measures and dependent variables were z-scored prior to modeling.

Prior to inference, assumptions of parametric tests were evaluated; when violations were detected, appropriate non-parametric alternatives were used. For repeated-measures ANOVAs, normality and outliers were assessed using the Shapiro-Wilk test; rank-based ANOVAs were applied when assumptions were violated, followed by Holm-corrected pairwise comparisons for significant effects. Linear regression models were fitted on standardized (z-scored) data, with 95% confidence intervals and p-values obtained using Wald $t$-distribution approximations. Model assumptions were assessed via residual normality (Shapiro-Wilk test), linearity (residual plots), homogeneity of variance (non-constant variance test), multicollinearity (variance inflation factor; VIF $< 5$), and outlier inspection. Linear mixed-effects models followed the same standardization and assumption-checking procedures. Competing linear models were compared against null models using Akaike Information Criterion (AIC), with the best-fitting model selected based on lowest AIC weight. Bayesian beta regression models were estimated using four chains of 2,000 iterations (1,000 warmup), yielding 4,000 post-warmup samples. Convergence and reliability were assessed using the Gelman–Rubin statistic (R-hat$<$1.05) and effective sample sizes (bulk and tail ESS$>$400). Model adequacy was further evaluated using posterior predictive checks, and only models meeting these criteria were reported.

<u>Computational Modelling</u>

*Hierarchical Bayesian Reinforcement learning models*

Learning parameters were estimated using hierarchical Bayesian reinforcement-learning models implemented in the *hBayesDM* package in R (54). Hierarchical estimation was chosen because it yields superior parameter recovery and statistical power relative to non-hierarchical approaches by jointly constraining group- and individual-level distributions (8, 55). For both experiments, multiple candidate models were fit (Table S5, S20) using four Markov chains with 4,000 iterations each (1,000 burn-in). Models were fit to trial-wise patch choices (1–3), gains (smiley emoji: 0/1), and losses (sad emoji: 0/–1). In Experiment 1, hierarchical priors were specified separately for anxiety group (high/low) and block (six blocks). In Experiment 2, priors were specified separately for session (pre-/post-sleep) and block (four blocks per session). Model convergence and reliability were assessed using R-hat (<1.05), effective sample size (ESS and tail-ESS), and Pareto-k diagnostics. Model comparison was based on Leave-One-Out Information Criterion (LOOIC), and the best-fitting model was selected as the one with the lowest LOOIC and no convergence or reliability violations (Table S4, S18). Additional model comparisons tested alternative prior structures, including pooled, group-only (anxiety or session), block-only, and group-by-block priors (Table S5, S19). In Experiment 1, a five-parameter model with separate reward and punishment learning rates, sensitivities, and a noise parameter provided the best fit. Incorporating both anxiety- and block-specific priors yielded the lowest LOOIC, indicating differential block-wise learning between anxiety groups. In Experiment 2, the best-fitting model additionally included a decay parameter, with session-and-block priors providing the best fit, indicating distinct learning across blocks and sleep sessions.

Model adequacy was evaluated using posterior predictive checks. Choices simulated from the winning model were compared with empirical behavior by correlating switch probabilities across trials; correlations exceeding 0.7 were taken as evidence of good model fit (8). The winning models showed strong correspondence with observed data (Pearson's correlation, Experiment 1: $r(46)=0.92$, $p<.001$; Experiment 2: pre-sleep: $r(38)=0.86$, $p<.05$; post-sleep: $r(38)=0.88$, $p<.05$). Group-level differences in parameters were assessed using 95% highest density intervals (HDIs) on hyperparameter differences, with effects considered reliable when HDIs did not include zero. In Experiment 1, post hoc analyses of individual-level parameter estimates were conducted using Bayesian mixed models (*brms*) to assess effects of stochasticity, volatility, and trait anxiety, including continuous symptom analyses with STAI-Y scores. In Experiment 2, mixed-effects regressions were conducted on individual-level parameter estimates to quantify sleep-related changes in learning. For each participant and block, the difference between post-sleep and pre-sleep parameter values (e.g., change in reward learning rate) was computed and used as the dependent variable. These difference scores indexed sleep-related modulation of model parameters and were analyzed to evaluate the joint effects of sleep (including N3 duration) and anxiety across all blocks.

To identify optimal parameter values that maximized rewards within each block, we simulated behavior using the best-fitting model across a broad parameter space. For each model parameter, nine values were selected spanning the empirical range of estimates; for learning rates, values ranged from 0.1 to 0.9. This resulted in 59,049 unique parameter combinations. For each parameter set, choices, gains, and losses were simulated across blocks. The top-performing 0.1% of simulated agents (50 agents per block) with the highest total rewards were identified, and their parameter values were averaged to define optimal parameters for that block. Individual deviation scores were then computed as the difference between participants' parameter estimates and the corresponding optimal values. Deviation scores were evaluated using Bayesian one-sample *t*-tests against a population mean of zero, with Bayesian correlations and tests conducted using *rstanarm* and *bayestestR* (56, 57). Effects were considered meaningful when 95% credible intervals excluded zero, and Bayes factors exceeded 1.

*Hierarchical Drift-Diffussion models*

Process-level modeling of the task behavior was performed using hierarchical drift-diffusion modeling was performed using HDDM package (v0.9.8) within a containerized Python environment (Docker), ensuring reproducible software dependencies. The drift-diffusion model comprises of four parameters estimated from trial-wise reaction times (RTs): drift rate ($v$), reflecting the speed of evidence accumulation towards an option; boundary separation ($a$), represents how much information is required to be accumulated to make a choice; starting-point bias ($z$); and non-decision time ($t$) captures sensory and motor processes unrelated to evidence accumulation (58). Choices were modeled using two boundaries: selecting the most rewarding patch versus any other patch (1/0). In Experiment 1, model parameters were allowed to vary as a function of the block and trait anxiety group (high/low). In Experiment 2, parameters varied by block and session (pre-/post-sleep). For each experiment, 10 candidate hierarchical models were fit, differing in prior structure (single pooled, block-only, anxiety/session-only, or anxiety/session×block) and parameter inclusion (full model or reduced models including subsets of parameters). All models were estimated using four chains with 4,000 samples each (1,000 burn-in). Model selection was based on convergence (R-hat<1.05) and Deviance Information Criterion (DIC), with the lowest DIC identifying the winning model.

In Experiment 1, the best-fitting model included block- and anxiety-dependent effects on drift rate and boundary separation (Table S9, S21). In Experiment 2, the winning model included session- and block-dependent effects on drift rate, boundary separation, and starting-point bias, with non-decision time varying by session only (Table S23). Model adequacy was assessed using posterior predictive checks. For each winning model, 500 datasets were simulated using parameters sampled from participants' posterior distributions (300 trials per simulation). Summary statistics from the empirical data fell within the 95% credible intervals of the simulated data, indicating good model fit. Convergence and reliability were further confirmed using effective sample size (ESS) diagnostics. Group-level differences were assessed by comparing posterior distributions of parameters, with <5% overlap taken as evidence of reliable differences ($q$<.05, 59, 60). For Experiment 2, individual-level parameter estimates were analyzed using mixed-effects regressions. Sleep-related change scores (post-sleep - pre-sleep; e.g., change in drift rate) were computed per block and participant and used as dependent variables to assess the effects of sleep (including N3 duration) and trait anxiety.

### Supplementary Text

Experiment 1: Effect of trait anxiety on punishment learning rates and other model parameters

Based on the group-level hyperparameter estimates: HTA lower punishment learning rate than LTA in medium stochasticity-fast volatility environment (95% HDI [1.52, 9.91]). This effect was again observed when examining individual punishment learning rates with HTA having overall lower punishment learning rate than LTA but similar in slow-volatility-high stochasticity environment for both categorical and continuous scale analysis (**Fig S3A, 3B,** Bayesian Beta Regression: HTA:β=-0.55, 95%CI [-0.77, -0.32]; HTA x High Stochasticity x Fast Volatility: β=-0.76, 95%CI [-1.11, -0.40], Table S11). However, punishment learning rates for both HTA and LTA participants were below optimal level (HTA: β=-0.10, 95%CI [-0.13, -0.07], BF=12100; LTA: β=-0.13, 95%CI [-0.21, -0.06], BF=34.41), indicating that punishment learning rates did not aid reward maximization on the task.

Effect of trait anxiety on other parameters:

a) *Reward sensitivity:* Both HTA and LTA were suboptimal. Focusing only on the slow-volatility-high stochasticity environment, we found that HTA had a significantly higher reward sensitivity while LTA had lower reward sensitivity than the optimal value (HTA:β=1.07, 95%CI [0.66, 1.45], BF = 1740; LTA: β=-6.50, 95% CI [-7.28, -5.72], BF =3.09e+12).
b) *Punishment sensitivity:* While, group-level hyperparameter estimates for punishment sensitivity were high in HTA in the medium stochasticity-fast volatility environment (95% HDI [1.52, 9.91**]**). However, both HTA and LTA were suboptimal. In the slow-volatility-high stochasticity environment both had significantly lower punishment sensitivity than the optimal value (HTA: β=-3.74, 95% CI [-5.70, -1.79], BF = 34.49; LTA: β=-3.18, 95% CI [-3.42, -2.93], BF = 4.46e+16).
c) *Noise:* Characterised as randomness in responding, both HTA and LTA were suboptimal, in slow-volatility-high stochasticity environment, HTA had significantly higher noise while LTA had lower noise than the optimal value (HTA: β=0.03, 95% CI [0.01, 0.04], BF = 17.08; LTA: β=-0.03, 95% CI [-0.03, 0.03], BF= 7.92e+35).

*For Evidence #2 Reward learning rate*: In slow volatility conditions for reward learning rates, HTA did not deviate from optimal values in low stochasticity environment (β= 0.003, 95% CI [-0.06, 0.07], BF = 0.224), while LTA showed no significant deviations from the optimal value (Low: β= 0.03, 95% CI [0.00, 0.06], BF = 1.36; Medium: β=0.03, 95% CI [-0.04, 0.09], BF = 0.345; High: β=0.02, 95% CI [-0.07, 0.11], BF = 0.251).

Thus, in this environment, HTA showed higher reward sensitivity and greater noise than LTA, while punishment learning rates did not differ between the groups. Notably, both groups exhibited lower-than-optimal punishment learning rates and sensitivities, contrary to expectations. This confirms that reward-focused parameters primarily drove HTA's reduced performance. Thus, our findings reveal a nuanced pattern of feedback processing in individuals with high trait anxiety (HTA), characterized primarily by a reduced punishment learning rate, particularly in volatile and highly stochastic environments. However, a critical exception emerged in the stable yet highly uncertain condition, where this learning difference from punishment disappeared. In this specific context, HTA individuals adopted a distinct and suboptimal strategy, displaying significantly higher reward sensitivity and greater decision noise along with previously mentioned higher reward learning rate. It is plausible that this strategic shift was amplified by the task design itself; with a final monetary incentive. Reward-focused outcomes were likely perceived as more salient and informative than punishments.

<u>Experiment 2: Repeated-measures ANOVA on total rewards.</u>
Here, high stochasticity and fast volatility lowered rewards (Stochasticity: $F_{1,312}$=143.86, $p$<.05,$\eta^2_p$=0.32; Volatility: $F_{1,312}$=8.64, $p$<.05,$\eta^2_p$=0.03; Post-hoc t-test: Low>High: $t_{159}$=6.71, $p$<.05, 95%CI [5.87, 7.83], $d$=0.94; Slow>Fast: $t_{159}$=2.64, $p$<.05, 95%CI [0.65, 2.64], $d$=0.26). But stochasticity and volatility also significantly interacted with each other ($F_{1,312}$=5.05, $p$<.05,$\eta^2_p$=0.02). Effect of volatility was seen only in low stochasticity condition where fast volatility lowered rewards than slow volatility (Simple main effect: Low: $F_{1,158}$=9.70, $p$<.05, $\eta^2_p$=0.06; High: $F_{1,158}$=0.37, $p$=.54, $\eta^2_p$=0.002; Post-hoc t-test: Slow>Fast: $t_{79}$=3.66, $p$<.05, 95%CI [1.32, 4.48], $d$=0.41). While, effect of stochasticity was seen in both volatility conditions where high stochasticity lowered rewards than low stochasticity (Simple main effect: Slow: $F_{1,158}$=84.03, $p$<.05, $\eta^2_p$=0.35; Fast: $F_{1,158}$=57.41, $p$<.05, $\eta^2_p$=0.27; Post-hoc t-test: Low-Slow > High-Slow: $t_{79}$=7.96, $p$<.05, 95%CI [6.13, 9.79], $d$=0.96; Low-Fast > High-Fast:$t_{79}$=5.45, $p$<.05, 95%CI [4.19, 6.71], $d$=0.96). Thus, higher stochasticity reduced reward accumulation under both slow and fast volatility, whereas changes in volatility had no additional effect when stochasticity was high.

<u>Experiment 2: Effect of other sleep variables on anxiety.</u>
Beyond N3 sleep, we preregistered hypotheses linking anxiety to other sleep parameters based on prior work. Specifically, we predicted that higher trait anxiety would be associated with fewer REM episodes and longer REM latency, whereas higher state anxiety would be associated with longer sleep onset latency (AsPredicted #128319, H1c i–ii; Horvath et al., 2016). Firstly, the REM episodes regression model violated assumptions of normality and had to be discarded. Trait anxiety was positively associated with longer REM latency (Linear Regression: β=0.48, 95%CI [0.11, 0.85], $t_{34}$=2.62, $p$<.05), an effect not observed for state anxiety or subjective sleep quality, indicating a trait-specific contribution to REM latency. However, we did not find state anxiety to affect sleep onset latency (Linear Regression corrected with Wald's test: β=0.05, $t_{34}$= 0.31, $p$=.8). We also hypothesized that lower morning anxiety would be related to greater power in the delta band during N3 sleep (AsPredicted #128319 H1b; Simon et al., 2020). As expected, we found a positive effect of dissipation in state anxiety with N3 delta bandpower (Linear Regression: β=0.65, 95%CI [0.15, 1.15], $t_{34}$= 2.66, $p$<.05).

<u>Experiment 2: Selecting best regression model for N3 sleep analysis on performance.</u>
Firstly, EEG-measured total sleep time correlated significantly with subjective sleep reports (Pearson's correlation: r(38) = 0.36, p<.05), confirming measurement consistency. We analyzed the post-minus-pre sleep reward difference using linear mixed-effects models and reported results from the best model which met all assumptions with the lowest AIC values (Table S17):

Increase in post sleep rewards ~ N3 + Trait Anxiety + DiffStateAnxiety + REM Latency + Stochasticity + Volatility + } *Main effects*
N3:Trait Anxiety + N3:REM Latency + N3:DiffStateAnxiety + N3:probability +N3:hazard_rate +
Trait Anxiety:DiffStateAnxiety + Trait Anxiety:Stochasticity + Trait Anxiety:Volatility +
DiffStateAnxiety:Stochasticity + DiffStateAnxiety :Volatility +
REM Latency:Trait Anxiety + REM Latency:Stochasticity + REM Latency:Volatility + } *Two-way interactions*
N3:Trait Anxiety:Stochasticity + N3:Trait Anxiety:Volatility +
REM Latency:Trait Anxiety:Stochasticity + REM Latency:Trait Anxiety:Volatility +
N3:REM Latency:Stochasticity + N3:REM Latency:Volatility + (1|participant) } *Three-way interactions*

We included REM latency, N3 delta-band power, and overnight dissipation of state anxiety as predictors because each was associated with a distinct preregistered hypothesis, allowing us to test whether N3 sleep duration uniquely accounted for effects in HTA. N3 Delta bandpower violated the assumption of collinearity possibly due to higher correlation with N3 sleep variable and was discarded from the final model. REM latency significantly interacted with trait anxiety ($\beta$=0.22, 95%CI [0.01, 0.44], $t_{149}$=2.04, $p$<.05), particularly with stochasticity ($\beta$=-0.26, 95%CI [-0.51, -0.02], $t_{120}$=-2.12, $p$<.05). REM latency also interacted with N3 sleep ($\beta$=-0.38, 95%CI [-0.71, -0.05], $t_{144}$=-2.28, $p$<.05) under stochasticity ($\beta$=0.41, 95%CI [0.05, 0.77], $t_{120}$=2.23, $p$<.05). With regards to the increase in identification of post-sleep most rewarding patch, REM latency also interacted with N3 sleep ($\beta$=-0.39, 95%CI [-0.74, -0.04], $t_{118}$=-2.21, $p$<.05). For both total rewards and identify most-rewarding patch: greater N3 sleep with lower REM latency helped to increase it post-sleep. However, the interactions of REM latency on trait anxiety did not extend to identification of the most rewarding patch, as well as REM latency x N3 interaction was not seen in post-sleep reward learning rates, suggesting that REM latency may not interact at a deeper level on performance i.e., this interaction might be due to chance or needs further systematic manipulation. Thus, we only report N3 x Trait anxiety interaction in the main text as this effect is consistently seen in all the results.

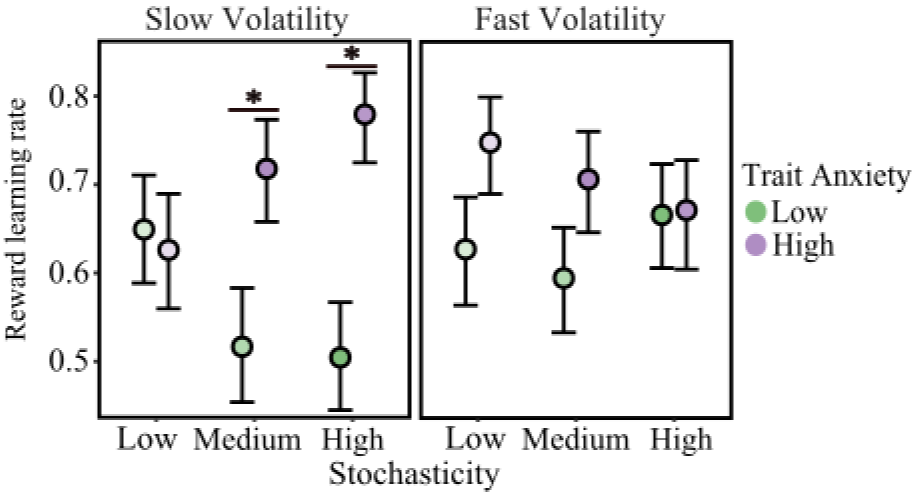


**Fig. S1. High trait anxiety and reward learning rates in slow-volatility, high-stochasticity environments.** Posterior means (± SEM) of reward learning rates show that when trait anxiety was treated categorically (HTA; STAI-T ≥ 45), reward learning rates did not differ between anxiety groups in fast-volatility environments. In contrast, under slow volatility, HTA individuals exhibited higher reward learning rates than LTA individuals as stochasticity increased. *Note*: Asterisks (✻) indicate statistically credible effects based on Bayesian inference.

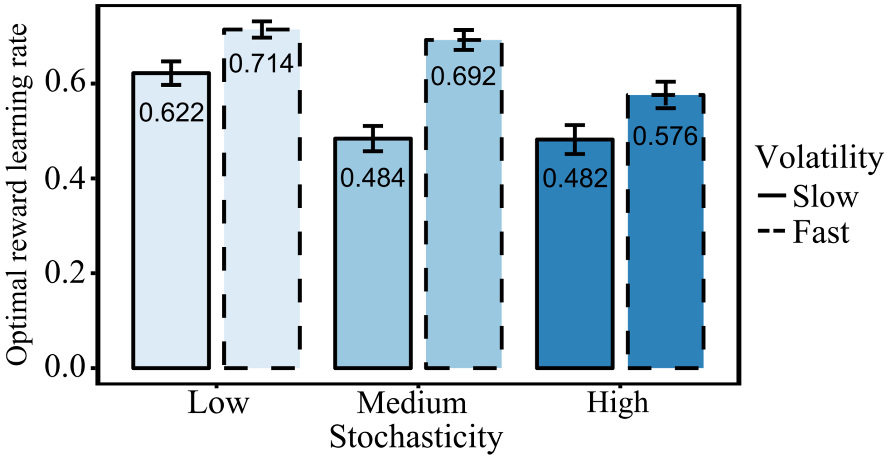


**Fig. S2. Evidence #2 Optimal reward learning rates across environments:** Higher stochasticity favors lower learning, while higher volatility favors higher learning for maximizing rewards. Bar plots contain mean (±SEM) optimal reward learning rate value for every environment.

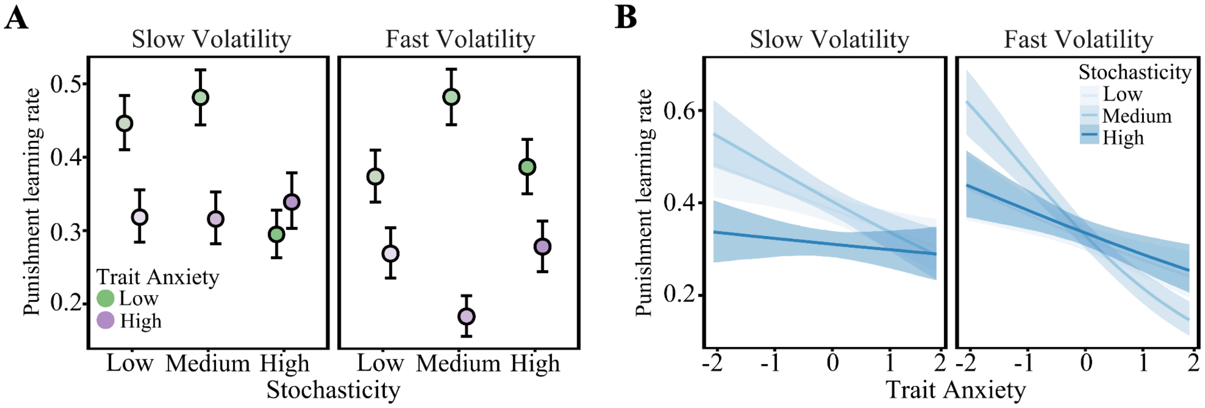


**Fig. S3. Effect of trait anxiety on punishment learning rates.** Punishment learning rates from Bayesian beta mixed regression models. **(A)** Posterior means (±SEM) of punishment learning rates for high- and low-trait-anxious (HTA, LTA) participants across volatility and stochasticity conditions. **(B)** Posterior estimates showing the relationship between continuous trait anxiety and punishment learning rates across volatility and stochasticity conditions. Both categorical and continuous trait anxiety scores showed lower punishment learning rates than LTA counterparts. In slow-volatility, high-stochasticity environment, rates were nearly identical across groups, suggesting that punishment learning did not drive performance in this environment.

**Table 1. Mixed effects ANOVA table on total rewards**

*Within-subject variables: stochasticity and volatility Between-subject variable: categorical trait anxiety (High trait anxious; STAI-T score ≥ 45). Significant differences are highlighted in bold where p<.05.*

| Predictor | | $df_{num}$ | $df_{den}$ | F | *p* | $\eta^2_p$ |
|---|---|---|---|---|---|---|
| Stochasticity | | 2 | 276 | 74.164 | **.000** | 0.350 |
| Volatility | | 1 | 276 | 7.692 | **.006** | 0.027 |
| Trait anxiety | | 1 | 276 | 4.076 | **.044** | 0.015 |
| Stochasticity x Volatility | | 2 | 276 | 0.472 | .624 | 0.003 |
| Stochasticity x Trait anxiety | | 2 | 276 | 0.538 | .585 | 0.004 |
| Volatility x Trait anxiety | | 1 | 276 | 4.866 | **.028** | 0.017 |
| Stochasticity x Volatility x Trait anxiety | | 2 | 276 | 1.093 | .337 | 0.008 |
| **Post-hoc pairwise comparisons** | | | **df** | **t** | ***p*** | *d* |
| Stochasticity | Low > Medium | | 95 | 7.74 | **.000** | 0.79 |
| | Low > High | | 95 | 13.67 | **.000** | 1.39 |
| | Medium > High | | 95 | 4.88 | **.000** | 0.5 |
| Volatility | Slow > Fast | | 144 | 3.00 | **.003** | 0.25 |
| Trait Anxiety | Slow volatility: HTA < LTA | | 129 | -2.3 | **.023** | -0.33 |
| | Fast volatility: HTA < LTA | | 129 | 0.11 | .911 | 0.03 |

*Note*: $df_{num}$ indicates degrees of freedom numerator, $df_{den}$ indicates degrees of freedom denominator, $\eta^2_p$ indicates partial eta-squared values i.e., the effect size, and *d* indicates Cohen's d effect size.

**Table 2. Between subject t-tests for every block/environment between HTA (group1) and LTA (group 2) individuals on total rewards.**

*To examine environment specific effects on total rewards. Significant differences are highlighted in bold where p<.05.*

| Block | df | t | *p* | 95% CI | *d* |
|---|---|---|---|---|---|
| Low Stochasticity- Slow Volatility | 41.05 | -0.53 | .598 | -4.36, 2.54 | -0.15 |
| Medium Stochasticity- Slow Volatility | 44.11 | -1.31 | .197 | -5.31, 1.12 | -0.38 |
| High Stochasticity- Slow Volatility | 36.78 | -2.61 | **.013** | -6.32, -0.79 | -0.77 |

| | | | | | |
|---|---|---|---|---|---|
| Low Stochasticity- Fast Volatility | 45.27 | 0.32 | .749 | -2.48, 3.43 | 0.09 |
| Medium Stochasticity- Fast Volatility | 44.63 | -0.44 | .660 | -2.81, 1.79 | -0.13 |
| High Stochasticity- Fast Volatility | 39.05 | 0.4 | .695 | -1.91, 2.84 | 0.12 |

*Note*: df indicates degrees of freedom, 95% CI indicates 95% confidence interval, and *d* indicates effect size using Cohen's *d*.

**Table 3. Effect on trialwise learning of the most rewarding patch.**
*Zero-one inflated beta (zoib) regression (fixed effects: stochasticity, volatility, trait anxiety; random effects: participant & trial) results in trial-wise learning of the most-rewarding patch. Model fitted with default log links for precision and logit for the mean, one-inflated and zero-inflated functions. Significant differences are highlighted (in bold) when 95% CI does not encompass zero.*

| | **Estimate (β)** | **SE** | **95% CI** |
|---|---|---|---|
| *Mean function* | | | |
| Intercept | 0.86 | 0.06 | 0.75 0.97 |
| Medium stochasticity | -0.49 | 0.02 | **-0.53 -0.44** |
| High Stochasticity | -0.57 | 0.02 | **-0.62 -0.53** |
| Fast volatility | -0.29 | 0.02 | **-0.33 -0.24** |
| HTA | -0.17 | 0.08 | -0.32 -0.01 |
| Medium stochasticity x Fast volatility | 0.05 | 0.03 | -0.01 0.11 |
| High stochasticity x Fast volatility | -0.1 | 0.03 | **-0.16 -0.04** |
| Medium stochasticity x HTA | -0.12 | 0.04 | **-0.18 -0.05** |
| High stochasticity x HTA | -0.19 | 0.03 | **-0.25 -0.13** |
| Fast volatility x HTA | 0.11 | 0.03 | **0.04 0.18** |
| Medium stochasticity x Fast volatility x HTA | 0.12 | 0.05 | **0.03 0.22** |
| High stochasticity x Fast volatility x HTA | 0.19 | 0.04 | **0.11 0.28** |
| *Precision function (phi)* | | | |
| Intercept | 2.45 | 0.13 | 2.19 2.7 |
| Medium stochasticity | -0.15 | 0.06 | **-0.27 -0.02** |
| High Stochasticity | 0.07 | 0.06 | -0.06 0.19 |

| | | | |
|---|---|---|---|
| Fast volatility | 0.29 | 0.07 | **0.15 0.42** |
| HTA | 0.2 | 0.18 | -0.18 0.54 |
| Medium stochasticity x Fast volatility | 0.25 | 0.09 | **0.07 0.42** |
| High stochasticity x Fast volatility | 0 | 0.09 | -0.17 0.18 |
| Medium stochasticity x HTA | -0.06 | 0.1 | -0.25 0.13 |
| High stochasticity x HTA | 0.12 | 0.09 | -0.06 0.31 |
| Fast volatility x HTA | -0.43 | 0.1 | **-0.63 -0.24** |
| Medium stochasticity x Fast volatility x HTA | -0.33 | 0.14 | **-0.61 -0.05** |
| High stochasticity x Fast volatility x HTA | -0.02 | 0.13 | -0.28 0.23 |
| *One-inflated function (coi)* | | | |
| Intercept | 3.64 | 0.74 | 2.26 5.17 |
| Medium stochasticity | -0.75 | 0.44 | -1.61 0.11 |
| High Stochasticity | -1.87 | 0.46 | **-2.77 -0.97** |
| Fast volatility | 0.11 | 0.47 | -0.81 1.03 |
| HTA | -0.38 | 0.89 | -2.11 1.35 |
| Medium stochasticity x Fast volatility | -0.67 | 0.65 | -2.01 0.6 |
| High stochasticity x Fast volatility | 0.76 | 0.67 | -0.61 2.02 |
| Medium stochasticity x HTA | -0.47 | 0.72 | -1.88 0.97 |
| High stochasticity x HTA | -0.03 | 0.74 | -1.45 1.45 |
| Fast volatility x HTA | -1.14 | 0.69 | -2.48 0.24 |
| Medium stochasticity x Fast volatility x HTA | 1.9 | 0.97 | -0.03 3.77 |
| High stochasticity x Fast volatility x HTA | 1.47 | 1.02 | -0.53 3.45 |
| *Zero-inflated function (zoi)* | | | |
| Intercept | -4.87 | 0.87 | -6.69 -3.27 |
| Medium stochasticity | -1.08 | 0.19 | **-1.46 -0.7** |
| High Stochasticity | -1.17 | 0.19 | **-1.55 -0.8** |

| | | | | |
|---|---|---|---|---|
| Fast volatility | -1.26 | 0.2 | **-1.64** | **-0.87** |
| HTA | -0.45 | 0.43 | -1.31 | 0.4 |
| Medium stochasticity x Fast volatility | 0.52 | 0.3 | -0.05 | 1.1 |
| High stochasticity x Fast volatility | 0.4 | 0.3 | -0.19 | 0.97 |
| Medium stochasticity x HTA | -0.61 | 0.31 | **-1.22** | **-0.01** |
| High stochasticity x HTA | -0.99 | 0.32 | **-1.63** | **-0.38** |
| Fast volatility x HTA | 0.58 | 0.29 | **0.01** | **1.12** |
| Medium stochasticity x Fast volatility x HTA | 0.87 | 0.45 | **0.02** | **1.74** |
| High stochasticity x Fast volatility x HTA | 1.06 | 0.45 | **0.16** | **1.93** |

**Table 4. Hierarchical reinforcement learning bayesian analysis model comparison**
*Model Comparisons with model specifications and LOOIC values. Highlighting (in bold) the best model with lowest LOOIC score.*

| Model | #Parameters | Parameters | | | | | | LOOIC |
|---|---|---|---|---|---|---|---|---|
| banditNarm_delta | 2 | Learning rate | Inverse temperature | | | | | 20312.99 |
| banditNarm_2par_lapse | 3 | Reward learning rate | Punishment learning rate | | | Lapse | | 25109.97 |
| banditNarm_4par | 4 | Reward learning rate | Punishment learning rate | Reward sensitivity | Punishment sensitivity | | | 17856.41 |
| banditNarm_lapse_decay | 6 | Reward learning rate | Punishment learning rate | Reward sensitivity | Punishment sensitivity | Lapse | Decay rate | 18012.77 |
| banditNarm_singleA_lapse | 4 | Learning rate | | Reward sensitivity | Punishment sensitivity | Lapse | | 18235.81 |
| banditNarm_lapse | 5 | Reward learning rate | Punishment learning rate | Reward sensitivity | Punishment sensitivity | Lapse | | **17851.88** |

**Table 5. Winning model prior fits**

*Model prior combinations of the best model. Selecting the best model with the best prior fits which have the lowest LOOIC values without any model convergence/reliability issues. [x]Denotes that the model did not converge, making the LOOIC values for the prior meaningless.*

| Priors | LOOIC |
|---|---|
| | banditNarm_lapse |
| Anxiety and Block Priors (12) | **17851.88** |
| Anxiety Priors (2) | 18379.17 |
| Block Priors (6) | 17755.31[x] |
| Single Prior (1) | 18354.88[x] |

**Table 6. Parameter estimates and group comparison on the winning model and prior combination.**

*Comparing parameter estimates of the best model between HTA v/s LTA for each block. In the HTA and LTA columns, the values represent the mean (standard deviation) of the average posterior estimates obtained from each individual. The 'between group HDI' column represents the lower and upper bounds of the 95% HDI. The intervals that do not encompass zero have been highlighted (in bold) as these are meaningful differences between the two groups. HDIs have been computed by taking the difference between HTA and LTA (HTA - LTA) for all parameters per block.*

| banditNarm_lapse | Block | HTA | LTA | Between group HDI |
|---|---|---|---|---|
| Reward learning rate | 80slow | 0.63 (0.03) | 0.65 (0.01) | -0.18 0.17 |
| | 70slow | 0.72 (0.02) | 0.52 (0.03) | **0.03 0.39** |
| | 60slow | 0.75 (0.04) | 0.51 (0.04) | **0.11 0.51** |
| | 80fast | 0.75 (0.01) | 0.63 (0.02) | -0.05 0.29 |
| | 70fast | 0.70 (0.03) | 0.60 (0.03) | -0.07 -0.33 |
| | 60fast | 0.66 (0.05) | 0.67 (0.01) | -0.19 0.21 |
| Punishment learning rate | 80slow | 0.32 (0.02) | 0.45 (0.00) | -0.41 0.11 |
| | 70slow | 0.31 (0.01) | 0.48 (0.01) | -0.35 0.02 |
| | 60slow | 0.34 (0.02) | 0.31 (0.03) | -0.21 0.26 |
| | 80fast | 0.27 (0.03) | 0.37 (0.01) | -0.27 0.06 |

| | | | | |
|---|---|---|---|---|
| | 70fast | 0.19 (0.02) | 0.48 (0.01) | **-0.51 -0.12** |
| | 60fast | 0.28 (0.01) | 0.39 (0.01) | -0.27 0.04 |
| Reward sensitivity | 80slow | 7.98 (0.69) | 7.08 (0.65) | -2.65 4.67 |
| | 70slow | 5.70 (0.62) | 6.31 (0.45) | -3.73 1.72 |
| | 60slow | 16.43 (0.19) | 8.90 (0.37) | -1.81 18.12 |
| | 80fast | 7.06 (0.61) | 7.79 (0.57) | -5.1 3.48 |
| | 70fast | 11.08 (0.61) | 8.61 (0.56) | -3.15 9.38 |
| | 60fast | 10.14 (0.76) | 7.08 (0.81) | -2.45 10.64 |
| Punishment sensitivity | 80slow | 4.00 (0.31) | 2.24 (0.21) | -1.24 4.71 |
| | 70slow | 3.52 (0.54) | 3.12 (0.41) | -1.9 2.62 |
| | 60slow | 5.06 (0.91) | 5.37 (0.12) | -5.95 3.72 |
| | 80fast | 5.47 (0.27) | 5.83 (0.68) | -3.51 3.6 |
| | 70fast | 8.20 (0.64) | 3.07 (0.57) | **1.52 9.91** |
| | 60fast | 4.98 (0.81) | 3.75 (0.61) | -2.03 3.94 |
| Lapse | 80slow | 0.02 (0.00) | 0.01 (0.00) | -0.03 0.03 |
| | 70slow | 0.02 (0.00) | 0.02 (0.00) | -0.04 0.05 |
| | 60slow | 0.08 (0.01) | 0.01 (0.00) | -0.003 0.11 |
| | 80fast | 0.02 (0.00) | 0.03 (0.00) | -0.05 0.04 |
| | 70fast | 0.02 (0.00) | 0.02 (0.00) | -0.03 0.04 |
| | 60fast | 0.03 (0.00) | 0.02 (0.00) | -0.04 0.06 |

**Table 7. Effect on reward learning rates**

*Reward learning rate results using bayesian beta regression for when trait anxiety was categorical and on a continuous scale. Significant differences are highlighted (in bold) when 95% CI does not encompass zero.*

| **Fixed effects** | ***Categorical Trait Anxiety*** | | | ***Continuous Trait Anxiety*** | | |
|---|---|---|---|---|---|---|
| | **β** | **SE** | **95% CI** | **β** | **SE** | **95% CI** |
| Intercept | 0.62 | 0.14 | **0.36 0.9** | 0.57 | 0.1 | 0.38 0.76 |

| | | | | | | | | |
|---|---|---|---|---|---|---|---|---|
| Medium Stochasticity | -0.55 | 0.15 | **-0.85** | **-0.26** | -0.11 | 0.11 | -0.33 | 0.11 |
| High Stochasticity | -0.6 | 0.15 | **-0.88** | **-0.32** | 0.00 | 0.11 | -0.22 | 0.23 |
| Fast Volatility | -0.1 | 0.15 | -0.39 | 0.2 | 0.2 | 0.11 | -0.04 | 0.41 |
| HTA | -0.1 | 0.2 | -0.5 | 0.28 | 0.07 | 0.09 | -0.12 | 0.25 |
| Medium Stochasticity:Fast Volatility | 0.41 | 0.21 | -0.01 | 0.82 | -0.06 | 0.16 | -0.37 | 0.26 |
| High Stochasticity:Fast Volatility | 0.77 | 0.21 | **0.36** | **1.18** | -0.08 | 0.16 | -0.39 | 0.24 |
| Medium Stochasticity:HTA | 0.97 | 0.22 | **0.53** | **1.42** | 0.36 | 0.12 | **0.13** | **0.59** |
| High Stochasticity:HTA | 1.34 | 0.23 | **0.9** | **1.79** | 0.62 | 0.12 | **0.39** | **0.85** |
| Fast Volatility:HTA | 0.67 | 0.23 | **0.21** | **1.11** | 0.17 | 0.12 | -0.06 | 0.4 |
| Medium Stochasticity:Fast Volatility:HTA | -1.04 | 0.32 | **-1.67** | **-0.41** | -0.41 | 0.16 | **-0.73** | **-0.1** |
| High Stochasticity:Fast Volatility:HTA | -1.89 | 0.32 | **-2.53** | **-1.25** | -0.83 | 0.17 | **-1.16** | **-0.5** |
| **Random effects** | | | | | | | | |
| Intercept: participant | 0.38 | 0.06 | 0.28 | 0.51 | 0.37 | 0.06 | 0.27 | 0.49 |
| **Family specific parameters** | | | | | | | | |
| Precision function | 14.11 | 1.28 | 11.71 | 16.79 | 13.90 | 1.25 | 11.56 | 16.47 |

**Table 8. Evidence #1 Comparing learning rates with rewards earned.**
*Comparing relationship between reward and punishment learning rates and total rewards earned in every block using bayesian correlations. Significant differences (when %ROPE is less than 1) are highlighted (in bold).*

| Parameter | Block | Median | 95% CI | Probability of Direction (pd) | %ROPE | Bayes factor (BF) |
|---|---|---|---|---|---|---|
| Reward learning rate | 80slow | -0.05 | -0.32, 0.20 | 65.50% | 28.03% | 0.348 |
| | 70slow | -0.23 | -0.47, 0.03 | 95.73% | 6.79% | 1.52 |
| | 60slow | -0.36 | -0.57, -0.09 | 99.35% | **0%** | 10.80 |
| | 80fast | -0.01 | -0.27, 0.27 | 52.20% | 28.29% | 0.325 |

| | | | | | | |
|---|---|---|---|---|---|---|
| | 70fast | 0.27 | -0.01, 0.50 | 97.30% | 3.61% | 2.11 |
| | 60fast | 0.19 | -0.07, 0.44 | 92.95% | 10.47% | 0.863 |
| Punishment learning rate | 80slow | -0.03 | -0.28, 0.25 | 56.85% | 28.89% | 0.328 |
| | 70slow | 0.11 | -0.16, 0.37 | 79.22% | 22.18% | 0.449 |
| | 60slow | -0.12 | -0.37, 0.17 | 81.15% | 20.53% | 0.473 |
| | 80fast | -0.13 | -0.38, 0.14 | 82.83% | 19.71% | 0.505 |
| | 70fast | 0.05 | -0.21, 0.32 | 63.98% | 28.95% | 0.351 |
| | 60fast | -0.16 | -0.41, 0.12 | 87.90% | 14.92% | 0.674 |

**Table 9. Evidence #3 Hierarchical drift-diffusion model comparisons**
*Model Comparisons with model specifications and DIC values. Highlighting (in bold) the best model i.e., with the lowest DIC score. Depends_on column denotes which parameters were fit on which priors. [x]Denotes that the model did not converge (R-hat value greater than 1.05), making the DIC values for the prior meaningless.*

| **Priors** | **Parameters depends_on** | | | | **DIC** |
|---|---|---|---|---|---|
| Anxiety * Block | $v$ : anxiety, block | $a$ : anxiety, block | $t$ : anxiety, block | $z$ : anxiety, block | 40076.67[x] |
| Anxiety*Block | $v$ : anxiety, block | $a$ : anxiety, block | $t$ : anxiety | $z$ : anxiety | **42040.86** |
| Anxiety*Block | $v$ : anxiety, block | $a$ : anxiety, block | $t$ : anxiety | $z$ : anxiety, block | 43379.99 |
| Anxiety*Block | $v$ : anxiety, block | $a$ : anxiety | $t$ : anxiety | $z$ : anxiety | 45524.81 |
| Block | $v$ : block | $a$ : block | $t$ : block | $z$ : block | 41417.28[x] |
| Block | $v$ : block | $a$ : block | $t$ | $z$ : block | 43376.46 |
| Block | $v$ : block | $a$ : block | $t$ | $z$ | 43519.07 |
| Block | $v$ : block | $a$ | $t$ | $z$ | 45519.17 |
| Anxiety | $v$ : anxiety | $a$ : anxiety | $t$ : anxiety | $z$ : anxiety | 46272.20 |
| Single | $v$ | $a$ | $t$ | $z$ | 46271.60 |

**Table 10. Evidence #3 Parameters estimates of drift-diffusion model**
*Comparing parameter estimates of the best model between HTA v/s LTA for each block by using their posterior distributions. In the HTA and LTA columns, the values represent the mean (standard deviation) of the average posterior estimates obtained from each individual*

| **Parameter** | **Block** | **HTA** | **LTA** | **Direction of comparison** | **% difference** | ***q*** |
|---|---|---|---|---|---|---|
| Drift-rate (*v*) | 80slow | 0.51 (0.07) | 0.58 (0.06) | HTA < LTA | 76.93% | 0.231 |
| | 70slow | 0.08 (0.07) | 0.29 (0.06) | | 98.97% | **0.010** |
| | 60slow | 0.03 (0.07) | 0.19 (0.06) | | 94.84% | **0.052** |
| | 80fast | 0.25 (0.07) | 0.26 (0.06) | | 55.04% | 0.450 |
| | 70fast | 0.07 (0.07) | 0.09 (0.06) | | 54.81% | 0.452 |
| | 60fast | -0.09 (0.07) | -0.04 (0.06) | | 71.06% | 0.289 |
| Boundary / threshold (*a*) | 80slow | 1.84 (0.09) | 1.93 (0.08) | HTA > LTA | 22.66% | 0.773 |
| | 70slow | 2.11 (0.09) | 2.11 (0.08) | | 51.32% | 0.487 |
| | 60slow | 1.99 (0.09) | 1.79 (0.08) | | 95.08% | **0.049** |
| | 80fast | 2.12 (0.09) | 2.11 (0.08) | | 52.74% | 0.473 |
| | 70fast | 1.77 (0.09) | 1.80 (0.08) | | 42.01% | 0.579 |
| | 60fast | 2.06 (0.09) | 1.88 (0.08) | | 92.10% | 0.079 |
| Non-decision time (*t*) | Overall | 1.92 (0.06) | 1.96 (0.05) | HTA < LTA | 70.99% | 0.290 |
| Bias (*z*) | Overall | 0.01 (0.02) | -0.04 (0.02) | HTA > LTA | 92.24% | 0.078 |

*Note:* % difference indicates the percentage of the difference between the two posterior distributions, q indicates the q-value i.e., percentage overlap with significant differences (<0.05) highlighted (in bold). Drift-rate 60slow q-value is near significance which is also highlighted.

**Table 11. Effect on punishment learning rates**

*Punishment learning rate results using bayesian beta regression for when trait anxiety was categorical and on a continuous scale.*

| **Fixed effects** | ***Categorical Trait Anxiety*** | | | ***Continuous Trait Anxiety*** | | |
|---|---|---|---|---|---|---|
| | **β** | **SE** | **95% CI** | **β** | **SE** | **95% CI** |
| Intercept | -0.22 | 0.08 | **-0.36 -0.06** | -0.47 | 0.06 | **-0.59 -0.34** |
| Medium Stochasticity | 0.14 | 0.08 | -0.02 0.3 | 0.07 | 0.07 | -0.07 0.21 |
| High Stochasticity | -0.66 | 0.09 | **-0.83 -0.49** | -0.33 | 0.07 | **-0.48 -0.19** |
| Fast Volatility | -0.3 | 0.08 | **-0.46 -0.14** | -0.27 | 0.07 | **-0.41 -0.13** |

| | | | | | | | | |
|---|---|---|---|---|---|---|---|---|
| HTA | -0.55 | 0.12 | **-0.77** | **-0.32** | -0.2 | 0.06 | **-0.32** | **-0.07** |
| Medium Stochasticity:Fast Volatility | 0.3 | 0.12 | **0.07** | **0.53** | -0.05 | 0.11 | -0.26 | 0.16 |
| High Stochasticity:Fast Volatility | 0.71 | 0.12 | **0.48** | **0.95** | 0.38 | 0.11 | **0.18** | **0.59** |
| Medium Stochasticity:HTA | -0.15 | 0.13 | -0.4 | 0.09 | -0.09 | 0.08 | -0.24 | 0.06 |
| High Stochasticity:HTA | 0.75 | 0.13 | **0.5** | **0.99** | 0.14 | 0.08 | -0.01 | 0.29 |
| Fast Volatility:HTA | 0.06 | 0.13 | -0.19 | 0.31 | -0.03 | 0.08 | -0.18 | 0.12 |
| Medium Stochasticity:Fast Volatility:HTA | -0.79 | 0.19 | **-1.15** | **-0.42** | -0.27 | 0.11 | **-0.47** | **-0.05** |
| High Stochasticity:Fast Volatility:HTA | -0.76 | 0.18 | **-1.11** | **-0.4** | -0.13 | 0.11 | -0.34 | 0.09 |
| **Random effects** | | | | | | | | |
| Intercept: participant | 0.24 | 0.03 | 0.19 | 0.32 | 0.27 | 0.04 | 0.20 | 0.35 |
| **Family specific parameters** | | | | | | | | |
| Precision function | 45.21 | 4.18 | 37.32 | 53.51 | 32.91 | 3.09 | 26.92 | 39.37 |

**Table 12. Comparing all sleep stages dissipation of state anxiety**
*Pearson's correlation between sleep stages and dissipation of state anxiety (pre-post sleep state). Only N3 sleep is significantly correlating with change in morning anxiety.*

| Sleep Stage | r | df | *p* |
|---|---|---|---|
| N1 | 0.17 | 38 | 0.30 |
| N2 | 0.03 | 38 | 0.84 |
| N3 | **0.36** | 38 | **0.02** |
| REM | 0.21 | 38 | 0.20 |

**Table 13. Effect of N3 sleep on dissipation in state anxiety**
*Linear regression results where dissipation in state anxiety was the dependent variable, with trait anxiety, state anxiety and SCI as independent variables.*

| Predictors | β | 95% CI | SE | t | *p* |
|---|---|---|---|---|---|
| Intercept | -0.03 | -0.32, 0.27 | 0.14 | -0.18 | .860 |

| | | | | | |
|---|---|---|---|---|---|
| N3 | 0.46 | 0.15, 0.77 | 0.15 | 3.01 | **.005** |
| Trait Anxiety | -0.23 | -0.54, 0.08 | 0.15 | -1.52 | .139 |
| SCI | -0.31 | -0.63, 0.07 | 0.16 | -1.99 | .055 |
| N3:Trait Anxiety | 0.43 | 0.06, 0.79 | 0.18 | 2.39 | **.023** |
| N3:SCI | -0.0007 | -0.37, 0.37 | 0.18 | -0.004 | .997 |
| Trait Anxiety:SCI | -0.03 | -0.41, 0.36 | 0.19 | -0.14 | .893 |
| N3:Trait Anxiety:SCI | -0.13 | -0.66, 0.39 | 0.26 | -0.53 | .599 |

**Table 14. Repeated-measures ANOVA table on total rewards**
*Within-subject variables: stochasticity, volatility and session. Significant differences are highlighted in bold where p<.05.*

| **Predictor** | | **$df_{num}$** | **$df_{den}$** | **F** | ***p*** | **$\eta^2_p$** |
|---|---|---|---|---|---|---|
| Stochasticity | | 1 | 312 | 143.86 | **.000** | 0.316 |
| Volatility | | 1 | 312 | 8.64 | **.004** | 0.027 |
| Session | | 1 | 312 | 8.91 | **.003** | 0.028 |
| Stochasticity:Volatility | | 1 | 312 | 5.05 | **.025** | 0.016 |
| Stochasticity:Session | | 1 | 312 | 0.25 | .615 | 0.001 |
| Volatility:Session | | 1 | 312 | 0.21 | .647 | 0.001 |
| Stochasticity:Volatility:Session | | 1 | 312 | 0.05 | .832 | 0.000 |
| **Post-hoc pairwise comparisons** | | **df** | **t** | ***p*** | *d* | |
| Stochasticity | Low > High | 159 | 6.71 | **.000** | 0.94 | |
| Volatility | Slow > Fast | 159 | 3.27 | **.001** | 0.26 | |
| Session | Post > Pre | 159 | 3.33 | **.001** | 0.26 | |
| Stochasticity:Volatility | Low Stochasticity: Slow > Fast | 79 | 3.66 | **.000** | 0.41 | |
| | High Stochasticity: Slow > Fast | 79 | -0.785 | .513 | 0.07 | |
| | Slow Volatility: Low > High | 79 | 7.96 | **.000** | 0.96 | |

| | | | | |
|---|---|---|---|---|
| Fast Volatility: Low > High | 79 | 5.45 | **.000** | 0.96 |

*Note*: $df_{num}$ indicates degrees of freedom numerator, $df_{den}$ indicates degrees of freedom denominator, $\eta^2_p$ indicates partial eta-squared values i.e., the effect size, and *d* indicates Cohen's d effect size.

**Table 15. Linear mixed-effect model comparison for sleep analysis**

*Model comparisons using AIC values to assess which variable combination is better at predicting the increase in post-sleep rewards. We used a combination of predictors: N3 , REM Latency, Trait Anxiety , DiffState Anxiety , Delta N3 , Stochasticity , Volatility. The best model with lowest AIC value is highlighted in bold.*

| **Predictors** | **AIC** |
|---|---|
| All variables but only main effects | 505.80 |
| All variables interaction i.e., 7th order | 509.80 |
| Without stochasticity & volatility interaction, 6th order | 516.12 |
| Without Delta N3, ie., 5th order | 503.28 |
| 4th order interaction | 491.5 |
| 3rd order interaction, without stochasticity and volatility interaction | **470.9** |
| 3rd order interaction but with stochasticity and volatility interactions | 472.76 |

**Table 16. Effect of N3 sleep on increase in post-sleep rewards**

*Linear mixed model (fixed effects: N3 time, REM Latency, Stochasticity, Volatility, Trait Anxiety, Morning State Anxiety; random effect: participant) results on increase in post-sleep rewards.*

| **Fixed effects** | **β** | **SE** | **95% CI** | ***df*** | ***t*** | ***p*** |
|---|---|---|---|---|---|---|
| Intercept | -0.1 | 0.13 | -0.36 0.16 | 140.91 | -0.78 | 0.435 |
| N3 | -0.09 | 0.16 | -0.4 0.22 | 148.74 | -0.58 | 0.561 |
| Trait Anxiety | -0.39 | 0.16 | -0.7 -0.09 | 141.41 | -2.53 | **0.013** |
| Dissipation in State Anxiety | 0.05 | 0.17 | -0.28 0.38 | 117.41 | 0.3 | 0.764 |
| REM Latency | -0.1 | 0.14 | -0.38 0.18 | 147.63 | -0.69 | 0.489 |
| High Stochasticity | -0.05 | 0.14 | -0.33 0.23 | 120 | -0.32 | 0.747 |
| Fast Volatility | 0.11 | 0.14 | -0.17 0.4 | 120 | 0.8 | 0.424 |
| N3:Trait Anxiety | 0.46 | 0.21 | 0.03 0.88 | 133.43 | 2.13 | **0.035** |

| | | | | | | | |
|---|---|---|---|---|---|---|---|
| N3:REM Latency | -0.38 | 0.17 | -0.71 | -0.05 | 143.87 | -2.28 | **0.024** |
| N3:Dissipation in State Anxiety | 0.08 | 0.08 | -0.09 | 0.25 | 40 | 0.95 | 0.35 |
| N3:High Stochasticity | 0.19 | 0.18 | -0.15 | 0.54 | 120 | 1.1 | 0.273 |
| N3:Fast Volatility | -0.07 | 0.18 | -0.42 | 0.27 | 120 | -0.42 | 0.676 |
| Trait Anxiety:REM Latency | 0.22 | 0.11 | 0.01 | 0.44 | 149.02 | 2.04 | **0.044** |
| Trait Anxiety:Dissipation in State Anxiety | -0.17 | 0.1 | -0.36 | 0.02 | 40 | -1.79 | 0.081 |
| Trait Anxiety:High Stochasticity | 0.13 | 0.17 | -0.2 | 0.47 | 120 | 0.78 | 0.437 |
| Trait Anxiety:Fast Volatility | 0.04 | 0.17 | -0.29 | 0.38 | 120 | 0.27 | 0.791 |
| Dissipation in State Anxiety:High Stochasticity | -0.26 | 0.16 | -0.58 | 0.07 | 120 | -1.57 | 0.119 |
| Dissipation in State Anxiety:Fast Volatility | 0.2 | 0.16 | -0.12 | 0.53 | 120 | 1.23 | 0.22 |
| REM Latency:High Stochasticity | 0.15 | 0.16 | -0.17 | 0.46 | 120 | 0.93 | 0.352 |
| REM Latency:Fast Volatility | -0.29 | 0.16 | -0.6 | 0.03 | 120 | -1.8 | 0.074 |
| N3:Trait Anxiety:High Stochasticity | -0.26 | 0.23 | -0.71 | 0.18 | 120 | -1.16 | 0.247 |
| N3:Trait Anxiety:Fast Volatility | -0.33 | 0.23 | -0.78 | 0.12 | 120 | -1.45 | 0.149 |
| Trait Anxiety:REM Latency:High Stochasticity | -0.26 | 0.12 | -0.51 | -0.02 | 120 | -2.12 | **0.036** |
| Trait Anxiety:REM Latency:Fast Volatility | 0.04 | 0.12 | -0.2 | 0.29 | 120 | 0.34 | 0.738 |
| N3:REM Latency:High Stochasticity | 0.41 | 0.18 | 0.05 | 0.77 | 120 | 2.23 | **0.028** |
| N3:REM Latency:Fast Volatility | -0.17 | 0.18 | -0.53 | 0.19 | 120 | -0.92 | 0.357 |
| **Random effects** | **Variance** | | | | **SD** | | |
| Intercept: participant | 0.03 | | | | 0.17 | | |

**Table 17. Effect of N3 sleep on increase in post-sleep identification of most rewarding patch**
*Linear mixed model (fixed effects: N3 time, REM Latency, Stochasticity, Volatility, Trait Anxiety, Morning State Anxiety ; random effect: participant) results on increase in identification of the most rewarding patch*

| Fixed effects | β | SE | 95% CI | df | t | p |
|---|---|---|---|---|---|---|
| Intercept | -0.17 | 0.14 | -0.44 0.11 | 114.23 | -1.22 | 0.225 |
| N3 | -0.02 | 0.17 | -0.35 0.31 | 124.38 | -0.12 | 0.902 |
| Trait Anxiety | -0.45 | 0.16 | -0.78 -0.13 | 114.82 | -2.74 | **0.007** |
| Dissipation in State Anxiety | 0.18 | 0.18 | -0.17 0.54 | 91.66 | 1.02 | 0.310 |
| REM Latency | -0.14 | 0.15 | -0.44 0.15 | 122.79 | -0.96 | 0.337 |
| High Stochasticity | -0.06 | 0.14 | -0.32 0.21 | 120.00 | -0.42 | 0.679 |
| Fast Volatility | 0.18 | 0.14 | -0.09 0.44 | 120.00 | 1.30 | 0.195 |
| N3:Trait Anxiety | 0.44 | 0.23 | -0.01 0.89 | 106.14 | 1.92 | **0.057** |
| N3:REM Latency | -0.39 | 0.18 | -0.74 -0.04 | 117.81 | -2.21 | **0.029** |
| N3:Dissipation in State Anxiety | 0.17 | 0.10 | -0.03 0.38 | 40.00 | 1.72 | 0.093 |
| N3:High Stochasticity | 0.00 | 0.17 | -0.33 0.34 | 120.00 | 0.03 | 0.980 |
| N3:Fast Volatility | 0.01 | 0.17 | -0.32 0.34 | 120.00 | 0.07 | 0.945 |
| Trait Anxiety:REM Latency | 0.20 | 0.12 | -0.03 0.43 | 124.78 | 1.72 | 0.087 |
| Trait Anxiety:Dissipation in State Anxiety | -0.21 | 0.11 | -0.44 0.02 | 40.00 | -1.86 | 0.070 |
| Trait Anxiety:High Stochasticity | 0.32 | 0.16 | 0.01 0.64 | 120.00 | 2.00 | **0.048** |
| Trait Anxiety:Fast Volatility | 0.13 | 0.16 | -0.19 0.44 | 120.00 | 0.78 | 0.439 |
| Dissipation in State Anxiety:High Stochasticity | -0.04 | 0.16 | -0.34 0.27 | 120.00 | -0.23 | 0.822 |
| Dissipation in State Anxiety:Fast Volatility | 0.08 | 0.16 | -0.23 0.38 | 120.00 | 0.49 | 0.623 |
| REM Latency:High Stochasticity | 0.09 | 0.15 | -0.21 0.39 | 120.00 | 0.58 | 0.564 |
| REM Latency:Fast Volatility | -0.17 | 0.15 | -0.47 0.13 | 120.00 | -1.14 | 0.256 |
| N3:Trait Anxiety:High Stochasticity | -0.10 | 0.22 | -0.53 0.32 | 120.00 | -0.48 | 0.631 |
| N3:Trait Anxiety:Fast Volatility | -0.49 | 0.22 | -0.92 -0.07 | 120.00 | -2.28 | **0.024** |
| Trait Anxiety:REM Latency:High Stochasticity | -0.15 | 0.12 | -0.38 0.08 | 120.00 | -1.27 | 0.207 |
| Trait Anxiety:REM Latency:Fast Volatility | -0.05 | 0.12 | -0.28 0.18 | 120.00 | -0.43 | 0.671 |

| | | | | | | | |
|---|---|---|---|---|---|---|---|
| N3:REM Latency:High Stochasticity | 0.34 | 0.17 | -0.01 | 0.68 | 120.00 | 1.93 | 0.056 |
| N3:REM Latency:Fast Volatility | 0.02 | 0.17 | -0.33 | 0.36 | 120.00 | 0.09 | 0.928 |
| **Random effects** | | | **Variance** | | | **SD** | |
| Intercept: participant | | | 0.15 | | | 0.38 | |

**Table 18. Hierarchical reinforcement learning bayesian analysis model comparison**
*Model Comparisons with model specifications and LOOIC values. Highlighting (in bold) the best model with lowest LOOIC score.*

| Model | #Parameters | Parameters | | | | | | LOOIC |
|---|---|---|---|---|---|---|---|---|
| banditNarm_delta | 2 | Learning rate | Inverse temperature | | | | | 23311.32 |
| banditNarm_2par_lapse | 3 | Reward learning rate | Punishment learning rate | | | Lapse | | 27713.21 |
| banditNarm_4par | 4 | Reward learning rate | Punishment learning rate | Reward sensitivity | Punishment sensitivity | | | 19850 |
| banditNarm_lapse_decay | 6 | Reward learning rate | Punishment learning rate | Reward sensitivity | Punishment sensitivity | Lapse | Decay rate | **19806.39** |
| banditNarm_singleA_lapse | 4 | Learning rate | | Reward sensitivity | Punishment sensitivity | Lapse | | 20153.49 |
| banditNarm_lapse | 5 | Reward learning rate | Punishment learning rate | Reward sensitivity | Punishment sensitivity | Lapse | | 19862.07 |

**Table 19. Winning model prior fits**
*Model prior combinations of the best model. Selecting the best model with the best prior fits which have the lowest LOOIC values without any model convergence/reliability issues. $^{x}$Denotes that the model did not converge, making the LOOIC values for the prior meaningless.*

| Priors | LOOIC |
|---|---|
| | banditNarm_lapse_decay |

| | |
|---|---|
| Session and Block Priors (8) | **19806.39** |
| Session Priors (2) | 20346.54 |
| Block Priors (4) | 20580.28 |
| Single Prior (1) | 20821.78 |

**Table 20. Effect of N3 sleep on post-sleep reward learning rates**

*Linear mixed model (fixed effects: N3 time, REM Latency, Stochasticity, Volatility, Trait Anxiety, Morning State Anxiety ; random effect: participant) results on increase in post-sleep reward learning rates. Learning rates from individual parameter estimates from the hierarchical Bayesian RL model.*

| **Fixed effects** | **β** | **SE** | **95% CI** | ***df*** | ***t*** | ***p*** |
|---|---|---|---|---|---|---|
| Intercept | 0.48 | 0.13 | 0.23 0.73 | 107.36 | 3.82 | **0.000** |
| N3 | 0 | 0.15 | -0.3 0.29 | 117.35 | -0.03 | 0.976 |
| Trait Anxiety | 0 | 0.15 | -0.3 0.29 | 107.93 | -0.02 | 0.985 |
| Dissipation in State Anxiety | 0.13 | 0.16 | -0.19 0.46 | 86.14 | 0.81 | 0.421 |
| REM Latency | -0.22 | 0.14 | -0.49 0.05 | 115.76 | -1.64 | 0.104 |
| High Stochasticity | -0.92 | 0.12 | -1.16 -0.69 | 120 | -7.77 | **0.000** |
| Fast Volatility | 0.05 | 0.12 | -0.19 0.28 | 120 | 0.42 | 0.677 |
| N3:Trait Anxiety | -0.45 | 0.21 | -0.86 -0.03 | 99.61 | -2.14 | **0.035** |
| N3:REM Latency | 0.18 | 0.16 | -0.14 0.49 | 110.84 | 1.11 | 0.270 |
| N3:Dissipation in State Anxiety | -0.04 | 0.09 | -0.23 0.15 | 40 | -0.45 | 0.658 |
| N3:High Stochasticity | -0.19 | 0.15 | -0.48 0.1 | 120 | -1.29 | 0.200 |
| N3:Fast Volatility | 0.05 | 0.15 | -0.24 0.35 | 120 | 0.37 | 0.712 |
| Trait Anxiety:REM Latency | -0.11 | 0.1 | -0.31 0.1 | 117.76 | -1.04 | 0.302 |
| Trait Anxiety:Dissipation in State Anxiety | 0.01 | 0.11 | -0.21 0.22 | 40 | 0.06 | 0.956 |
| Trait Anxiety:High Stochasticity | -0.07 | 0.14 | -0.35 0.21 | 120 | -0.5 | 0.621 |
| Trait Anxiety:Fast Volatility | 0.07 | 0.14 | -0.21 0.35 | 120 | 0.49 | 0.625 |
| Dissipation in State Anxiety:High | -0.01 | 0.14 | -0.28 0.26 | 120 | -0.11 | 0.915 |

| | | | | | | | |
|---|---|---|---|---|---|---|---|
| Stochasticity | | | | | | | |
| Dissipation in State Anxiety:Fast Volatility | -0.04 | 0.14 | -0.31 | 0.23 | 120 | -0.32 | 0.749 |
| REM Latency:High Stochasticity | 0.01 | 0.13 | -0.25 | 0.28 | 120 | 0.11 | 0.911 |
| REM Latency:Fast Volatility | 0.03 | 0.13 | -0.23 | 0.29 | 120 | 0.24 | 0.813 |
| N3:Trait Anxiety:High Stochasticity | -0.08 | 0.19 | -0.45 | 0.29 | 120 | -0.42 | 0.678 |
| N3:Trait Anxiety:Fast Volatility | 0.26 | 0.19 | -0.12 | 0.63 | 120 | 1.36 | 0.178 |
| Trait Anxiety:REM Latency:High Stochasticity | 0.13 | 0.1 | -0.07 | 0.34 | 120 | 1.3 | 0.198 |
| Trait Anxiety:REM Latency:Fast Volatility | 0.06 | 0.1 | -0.15 | 0.26 | 120 | 0.57 | 0.570 |
| N3:REM Latency:High Stochasticity | -0.11 | 0.15 | -0.42 | 0.19 | 120 | -0.74 | 0.46 |
| N3:REM Latency:Fast Volatility | -0.2 | 0.15 | -0.5 | 0.1 | 120 | -1.3 | 0.196 |
| **Random effects** | **Variance** | | | | **SD** | | |
| Intercept: participant | 0.14 | | | | 0.38 | | |

**Table 21. Hierarchical drift-diffusion model comparisons**

*Model Comparisons with model specifications and DIC values. Highlighting (in bold) the best model i.e., with the lowest DIC score. Depends_on column denotes which parameters were fit on which priors. [x]Denotes that the model did not converge (R-hat value greater than 1.05), making the DIC values for the prior meaningless.*

| **Priors** | **Parameters depends_on** | | | | **DIC** |
|---|---|---|---|---|---|
| Session* Block | $v$ : session, block | $a$ : session, block | $t$ : session, block | $z$ : session, block | 43152.91[x] |
| Session*Block | $v$ : session, block | $a$ : session, block | $t$ : session | $z$ : session, block | **44878.01** |
| Session*Block | $v$ : session, block | $a$ : session, block | $t$ : session | $z$ : session | 44902.26 |
| Session*Block | $v$ : session, block | $a$ : session | $t$ : session | $z$ : session | 46123.33 |
| Block | $v$ : block | $a$ : block | $t$ : block | $z$ : block | 47107.52 |
| Block | $v$ : block | $a$ : block | $t$ | $z$ : block | 48041.03 |
| Block | $v$ : block | $a$ : block | $t$ | $z$ | 48089.02 |

| | | | | | |
|---|---|---|---|---|---|
| Block | *v* : block | *a* | *t* | *z* | 48581.39 |
| Session | *v* : session | *a* : session | *t* : session | *z* : session | 46700.83 |
| Single | *v* | *a* | *t* | *z* | 49100.68 |

**Table 22. Effect of N3 sleep on post-sleep drift-rates**
*Linear mixed model (fixed effects: N3 time, REM Latency, Stochasticity, Volatility, Trait Anxiety, Dissipation in State Anxiety; random effect: participant) results on increase in post-sleep drift-rates.*

| **Fixed effects** | **β** | **SE** | **95% CI** | ***df*** | ***t*** | ***p*** |
|---|---|---|---|---|---|---|
| Intercept | -0.05 | 0.14 | -0.32 0.22 | 120.91 | -0.34 | 0.732 |
| N3 | 0.02 | 0.16 | -0.3 0.34 | 130.95 | 0.1 | 0.922 |
| Trait Anxiety | -0.52 | 0.16 | -0.84 -0.2 | 121.5 | -3.25 | **0.002** |
| Dissipation in State Anxiety | 0.06 | 0.17 | -0.29 0.41 | 97.35 | 0.34 | 0.734 |
| REM Latency | -0.13 | 0.15 | -0.42 0.16 | 129.4 | -0.88 | 0.382 |
| High Stochasticity | -0.24 | 0.14 | -0.51 0.03 | 120 | -1.77 | **0.079** |
| Fast Volatility | 0.12 | 0.14 | -0.15 0.39 | 120 | 0.89 | 0.373 |
| N3:Trait Anxiety | 0.48 | 0.22 | 0.03 0.92 | 112.64 | 2.13 | **0.035** |
| N3:REM Latency | -0.38 | 0.17 | -0.72 -0.04 | 124.49 | -2.21 | **0.029** |
| N3:Dissipation in State Anxiety | 0.2 | 0.1 | 0.01 0.39 | 40 | 2.06 | **0.046** |
| N3:High Stochasticity | -0.02 | 0.17 | -0.35 0.31 | 120 | -0.12 | 0.906 |
| N3:Fast Volatility | -0.04 | 0.17 | -0.38 0.29 | 120 | -0.26 | 0.798 |
| Trait Anxiety:REM Latency | 0.18 | 0.11 | -0.04 0.4 | 131.34 | 1.59 | 0.114 |
| Trait Anxiety:Dissipation in State Anxiety | -0.13 | 0.11 | -0.35 0.08 | 40 | -1.23 | 0.226 |
| Trait Anxiety:High Stochasticity | 0.4 | 0.16 | 0.07 0.72 | 120 | 2.44 | **0.016** |
| Trait Anxiety:Fast Volatility | 0.19 | 0.16 | -0.13 0.51 | 120 | 1.15 | 0.253 |
| Dissipation in State Anxiety:High Stochasticity | -0.03 | 0.16 | -0.34 0.28 | 120 | -0.17 | 0.869 |
| Dissipation in State Anxiety:Fast Volatility | 0.12 | 0.16 | -0.19 0.43 | 120 | 0.77 | 0.444 |
| REM Latency:High Stochasticity | 0.09 | 0.15 | -0.22 0.39 | 120 | 0.57 | 0.572 |

| | | | | | | | |
|---|---|---|---|---|---|---|---|
| REM Latency:Fast Volatility | -0.21 | 0.15 | -0.51 | 0.09 | 120 | -1.37 | 0.173 |
| N3:Trait Anxiety:High Stochasticity | -0.22 | 0.22 | -0.65 | 0.2 | 120 | -1.03 | 0.304 |
| N3:Trait Anxiety:Fast Volatility | -0.51 | 0.22 | -0.93 | -0.08 | 120 | -2.34 | **0.021** |
| Trait Anxiety:REM Latency:High Stochasticity | -0.11 | 0.12 | -0.35 | 0.12 | 120 | -0.95 | 0.347 |
| Trait Anxiety:REM Latency:Fast Volatility | -0.05 | 0.12 | -0.29 | 0.18 | 120 | -0.46 | 0.644 |
| N3:REM Latency:High Stochasticity | 0.4 | 0.17 | 0.05 | 0.74 | 120 | 2.26 | **0.025** |
| N3:REM Latency:Fast Volatility | 0.02 | 0.17 | -0.33 | 0.36 | 120 | 0.1 | 0.922 |
| **Random effects** | **Variance** | | | | **SD** | | |
| Intercept: participant | 0.11 | | | | 0.34 | | |

**Table 23. Average sleep statistics**
*Sleep statistics of participants i.e., mean (standard deviation) of the time spent / percentage in each stage and architecture.*

| **N = 40** | **Time (min)** | **% of Total sleep time** |
|---|---|---|
| Total sleep time (TST) | 383.31 (112.18) | |
| Sleep onset latency (SOL) | 31.01 (26.87) | |
| Wake after sleep onset (WASO) | 108.33 (86.42) | |
| N1 | 29.09 (20.80) | 6.97 (4.28) |
| N2 | 196.05 (49.03) | 52.82 (10.04) |
| N3 | 61.73 (27.56) | 16.62 (8.43) |
| REM | 96.45 (47.55) | 23.59 (8.65) |
| REM Latency | 120.79 (52.66) | |
